# A Hippocampal-parietal Network for Reference Frame Coordination

**DOI:** 10.1101/2024.08.21.609019

**Authors:** Yicheng Zheng, Xinyu Zhou, Shawn C. Moseley, Sydney M. Ragsdale, Leslie J. Alday, Wei Wu, Aaron A. Wilber

## Abstract

Navigating space and forming memories based on spatial experience are crucial for survival, including storing memories in an allocentric (map-like) framework and conversion into body-centered action. The hippocampus and parietal cortex (PC) comprise a network for coordinating these reference frames, though the mechanism remains unclear. We used a task requiring remembering previous spatial locations to make correct future action and observed that hippocampus can encode the allocentric place, while PC encodes upcoming actions and relays this to hippocampus. Transformation from location to action unfolds gradually, with ‘Came From’ signals diminishing and future action representations strengthening. PC sometimes encodes previous spatial locations in a route-based reference frame and conveys this to hippocampus. The signal for the future location appears first in PC, and then in hippocampus, in the form of an egocentric direction of future goal locations, suggesting egocentric encoding recently observed in hippocampus may originate in PC (or another “upstream” structure). Bidirectional signaling suggests a coordinated mechanism for integrating map-like, route-centered, and person-centered spatial reference frames at the network level during navigation.

## Introduction

Navigation is a fundamental cognitive process that allows organisms to move through their environment effectively. Two frames of reference are most frequently considered for understanding how we map out and navigate our surroundings: allocentric (world-centered) and egocentric (self-centered). The coordination between these two frameworks is critical for understanding how animals, including humans, interpret and interact with their surroundings. There are a number of computational and theoretical studies for how coordination and transformation between these reference frames occurs^1–5^. This transformation between allocentric and egocentric spatial representations is thought to involve multiple brain regions, including the hippocampus, parahippocampal regions, and anterior thalamic nucleus, where spatial representations are more frequently allocentric; sensory and motor cortex, where representations are more frequently egocentric; and retrosplenial cortex and parietal cortex (PC), where mixed allocentric and egocentric encoding has been observed^3,6^.

The hippocampus has long been implicated in allocentric spatial processing, a view supported by extensive research indicating its role in forming and retrieving allocentric spatial representations of the environment^7^. Place cells in the hippocampus encode specific locations in an environment, providing a neural basis for the allocentric framework^8^. In contrast, the PC is traditionally associated with egocentric spatial processing and integrates sensory information to produce a representation of the spatial relationships relative to the body. It also contains allocentric head direction (HD) cells and cells conjunctive for egocentric goal locations and HD^9,10^.

Despite these well-established roles, recent evidence suggests that a number of brain regions, including the hippocampus and even sensory cortices, might also encode information about ‘other’ spatial frameworks^11–16^, challenging the traditional dichotomy. For instance, hippocampal neurons were shown to be able to exhibit egocentric tuning^17–19^. These findings imply there is potential for a more integrated and dynamic relationship between these brain regions in spatial processing.

Understanding the interplay between the hippocampus and PC in coordinating between allocentric and egocentric information is crucial for elucidating the neural mechanisms underlying navigation in contexts that require rapid integration across reference frames, such as when biking or driving, where quick decisions must be made at an intersection. The hippocampus and PC are interconnected through various pathways and are frequently found to functionally interact^20^, allowing for the exchange of spatial information across this extended brain network which may serve to coordinate between allocentric and egocentric perspectives ^21^.

Here, we investigated how the hippocampus and PC coordinate between encoding of a previously visited spatial location and a future egocentric goal location during a complex spatial sequence task. This task requires rats to remember a prior spatial location and then generate an appropriate future action while traversing a common route. Our results show that both the hippocampus and PC encode and exchange information about the prior and future spatial locations, supporting the notion of a bidirectional coordination between allocentric and egocentric perspectives.

These findings extend our understanding of spatial navigation by highlighting the dynamic and flexible nature of spatial representations in the brain and the extent of the brain network which coordinates this information. They suggest that rather than operating in isolation, the hippocampus and PC, despite multiple synapses in between, interact closely to support the coordination between allocentric, route-centered, and egocentric perspectives for effective navigation.

## Results

Rats were trained to navigate a complex sequence of spatial locations on an open circular arena as described previously^22–24^. Among 32 possible spatial locations equally distributed around the perimeter of the large circular arena, five locations were arranged as an 8-item sequence (1-2-3-4-1-2-3-5-). Rats were trained to run through the sequence of locations with rewards at each location (Fig. 1a). Thus, during the 1-2-3 segment each rat had to remember its previous location (*came from* spatial location 4 vs. 5) to determine the next move (*go to* zone 4 vs. 5; Fig. 1 b). If they failed to reach the correct zone within 18 s, a light cue for the next rewarding zone would be activated to guide the rat to the correct location, and the trial would be counted as an error. Once well trained, rats accurately followed the sequence, making up to 90% correct choices (Fig. 1c).

**Fig. 1:**
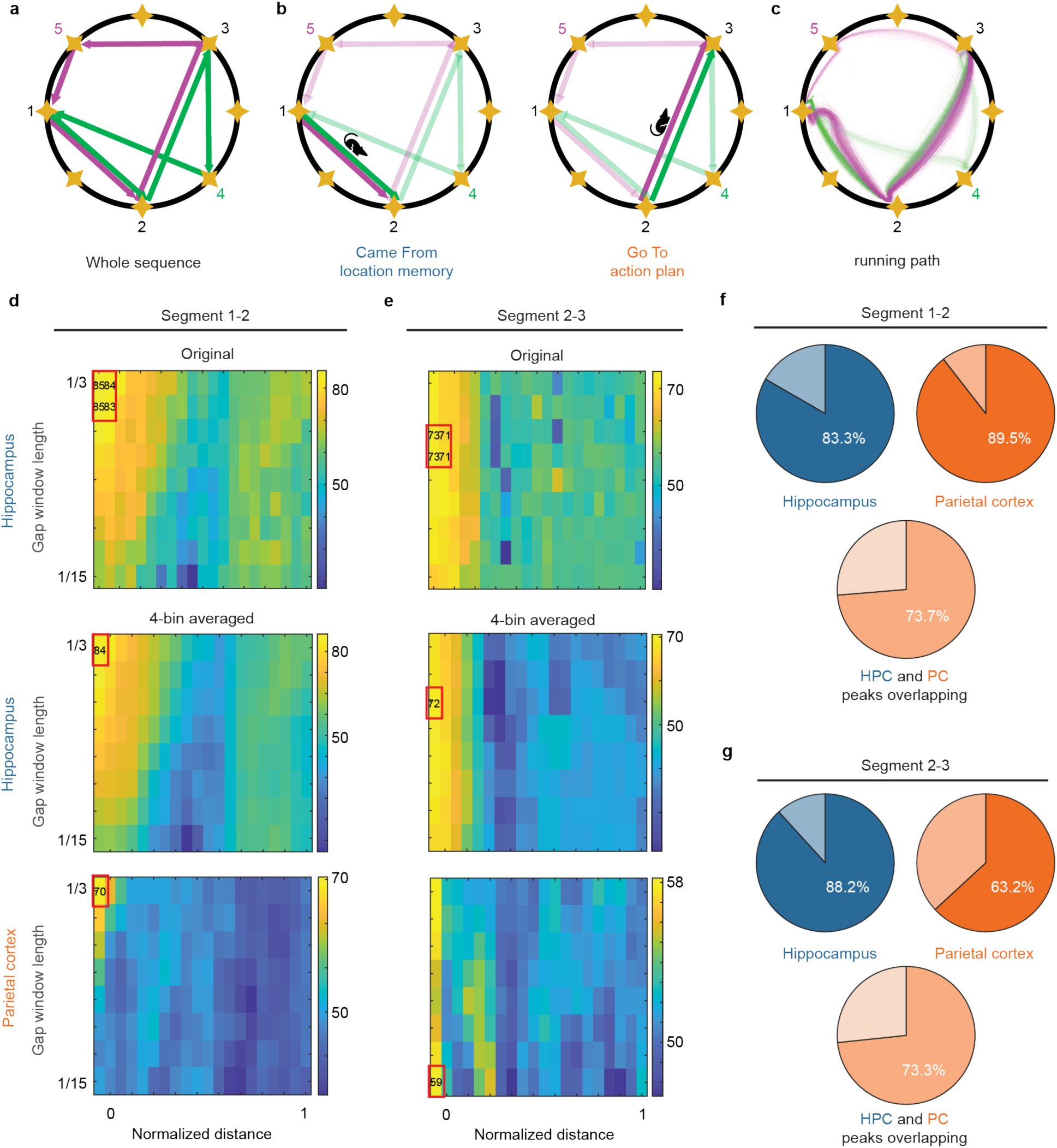
‘Came From’ and ‘Go To’ Information Can Be Decoded from Hippocampus and PC but Peaks from Different Brain Regions Occur at the Same Place on the Maze. **a**, Schematic of the complex sequence task. Path 1-2-3-4 is marked in green, and 1-2-3-5 is marked in magenta. **b**, To navigate the spatial sequence, the rat must remember where it “Came From” during segment 1-2 (*left*) and subsequently use this information to generate a plan for where it will ‘Go To’ during segment 2-3 (*right*). **c**, Example of the running path during one behavior session. **d, e**, Examples of spatial decoding matrix in segment 1-2 (**d**) and 2-3 (**e**). *Top*: Original decoding matrix for hippocampal single cells. *Middle*: Same as *top*, except that every set of 4 adjacent bins from the *top* was averaged to generate this 4-bin average map to extract the peak in the decoding matrix. *Bottom*: Same as *middle*, but for PC MUA. **f, g**, For segment 1-2 (**f**) and segment 2-3 (**g**), proportions are shown for the position of spatial decoding. *Top*. Proportion of sessions for which the spatial decoding peak is at the beginning of the segment. *Bottom*. Proportion of sessions for which the PC and hippocampus peaks overlap.

### ‘Came From’ information is apparent in both hippocampus and PC at the same spatial location on the 1-2 segment

We conducted neural decoding based on spatial position on segment 1-2 and 2-3. Spikes on actual running path were projected onto the straight lines directly connecting two rewarding zones. Our previous work showed modular organization of encoding in PC^23^. Thus, we initially analyzed multi-unit activity (MUA) and single units. This approach yielded encoding of many variables of interest in the MUA, so we focused on MUA for PC data. Thus, we recorded hippocampal single units and PC MUA (and some PC single units). Each segment was divided into 10 bins, and dìerent window sizes (i.e., larger windows included more spatial bins) were applied from the start point of each bin forward in time. This process generated a decoding matrix showing prediction accuracies for one behavior session as a function with position on the segment and decoding window size (Fig. 1d *Top*). To extract a peak in the decoding parameter space, each set of 4 adjacent bins were averaged to generate a 4-bin average decoding matrix (Fig. 1d *Middle*). The best prediction accuracy from this matrix was used to represent the decoding accuracy for the current behavior session (see Methods). Note that because we used a 4-bin average, the peak accuracy was reduced slightly compared to the original decoding matrix. We employed this approach because the peak in the decoding space varied particularly across time (as described below) between data sets and rats. Thus, extracting the peak allowed us to best capture this neural signal. Finally, the spatial locations for the PC and hippocampus peaks were confined to the beginning (within the first 3 spatial bins, a span of approximately 30 cm, of the 1-2 segment) for most data sets (Fig. 1f *Top*). These peaks were overlapping for most data sets (Fig. 1d *Middle and Bottom* and Fig. 1f *Bottom*).

### ‘Go To’ information is also apparent in both hippocampus and PC at the same spatial location on the 2-3 segment

Furthermore, we conducted ‘Go To’ decoding during segment 2-3 using the same decoding approach as described above, except that a model was built for the future goal location (zone 4 vs. 5). Similar to what we observed with ‘Came From’ decoding, the decoding peaks from hippocampal and PC data were again confined to the beginning of the segment for most data sets (Fig. 1g Top). As a result, PC and HPC decoding peaks overlapped in spatial location for most of the behavior sessions on segment 2-3 (Fig. 1e *Middle* and *Bottom* and Fig. 1g *Bottom*). Thus, we next assessed the temporal relationship between ‘Came From’ and ‘Go To’ signals in PC and hippocampus to see if there were temporal sequences within this spatial location.

### ‘Came From’ and ‘Go To’ signals are apparent in hippocampus and PC at different times

We next conducted neural decoding based on time. However, the running duration of each segment was normalized since rats tended to run more slowly near the end than at the beginning of each behavior session (Extended Fig. 1a). Thus, we normalized the time for running each segment by dividing each value by 5 s (longer than most running durations). Then, segments 1-2 and 2-3 were divided into 10 time bins, and a range of decoding window sizes were applied. As with spatial decoding, from the original decoding matrix (Fig. 2a, b *Top*), each 4 adjacent bins were averaged to generate a 4-bin averaged decoding matrix (Fig. 2a, b *Middle and Bottom*). The best prediction accuracy from this matrix was again used to represent the decoding performance for the current behavior session. To account for including multiple data sets from each rat, we used a mixed-effects model that accounted for rat identity to test significance between dìerent categories (Methods). We found that unlike spatial decoding, hippocampus and PC temporal decoding peaks did not overlap but were separated in time for both segments 1-2 and 2-3 (Fig. 2a, b). Moreover, we found that ‘Go To’ decoding peaks from PC MUA typically occurred before hippocampal peaks in segment 2-3 (t_(21)_=2.74, p=0.013; Fig. 2d *Right*), suggesting that ‘Go To’ information is conveyed from PC to hippocampus. However, we did not see a clear temporal order in segment 1-2, as decoding peaks from hippocampal single units occurred before PC peaks for some data sets and after PC peaks for other data sets (t_(21)_=0.35, p=0.73; Fig. 2d *Left*), suggesting a more complicated information flow for information about where the rat had come from.

**Fig. 2:**
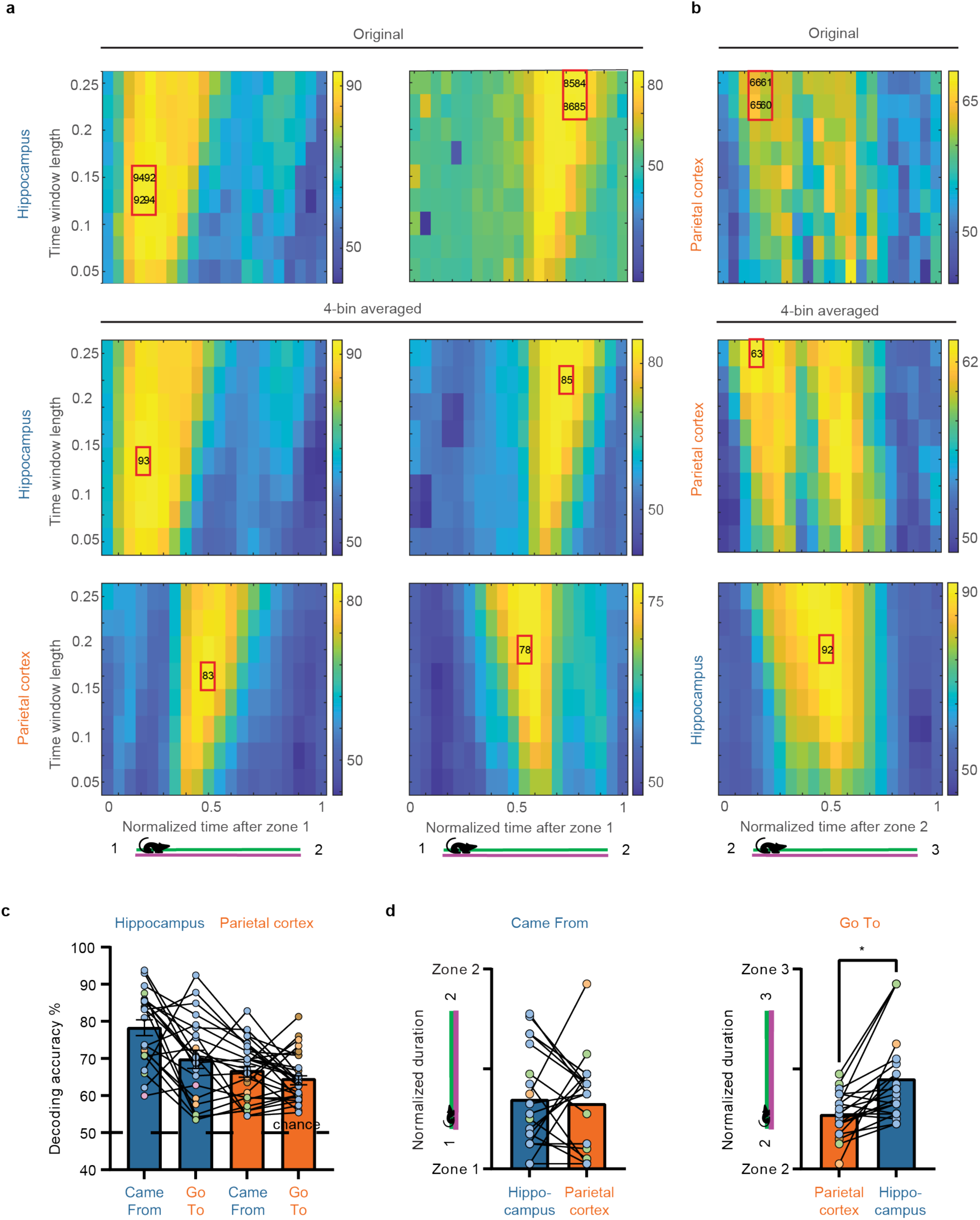
Temporal Decoding Reveals Sequential Patterns Across Hippocampus and PC of ‘Came From’ and ‘Go To’ Information. **a**, Examples of temporal decoding matrices for ‘Came From’ decoding during segment 1-2. *Top*: Original hippocampal decoding matrix. Horizontal axis is normalized time, and vertical axis is window size (proportion of normalized segment time). *Middle*: Each set of 4 adjacent bins is averaged to produce an average decoding matrix so that the peak in the parameter space can be extracted. *Bottom*: Averaged decoding matrix for PC. Color bar: Decoding accuracy (percent). **b**, Examples of temporal decoding matrices for ‘Go To’ decoding during segment 2-3. *Top*: Original PC decoding matrix. *Middle*: PC 4-bin averaged decoding matrix. *Bottom*: Hippocampus 4-bin averaged decoding matrix. **c**, Decoding accuracy varied across ‘Came From’ and ‘Go To’ decoding in hippocampus and PC. **d**, Temporal order of decoding peaks for ‘Came From’ decoding in segment 1-2 (*left*) and ‘Go To’ decoding in segment 2-3 (*right*). \**P* < 0.05; \*\*\**P* < 0.001.

Next, we compared accuracy for ‘Came From’ decoding in segment 1-2 and ‘Go To’ decoding in segment 2-3 in both hippocampus and PC, and found that the prediction accuracies were all well above chance, suggesting that both hippocampus and PC contained information about ‘Came From’ and ‘Go To’ (Fig. 2c). To compare accuracies between brain regions (hippocampus and PC) and decoding types (‘Came From’ and ‘Go To’), we used a mixed-effects model (see Methods), and found ‘Came From’ decoding accuracy is significantly higher than ‘Go To’ (F_(1)_=15.1; p<0.001). We also found that hippocampus decoding accuracy was significantly higher than PC (F_(1)_=67.0; p<0.001), however it is harder to know what to make of this “apples to oranges” comparison for MUA vs. single units. Finally, decoding category did not vary significantly across brain region (i.e., non-significant interaction; F_(1)_=0.86; p=0.36).

Finally, we compared temporal and spatial decoding accuracy for ‘Came From’ decoding during the 1-2 segment and ‘Go To’ decoding during the 2-3 segment for both hippocampus and PC, and found that for hippocampus, decoding did not vary significantly across methods (spatial vs. temporal) and types (‘Came From’ vs. ‘Go To’; F_(1)_=3.5, p=0.07). However, temporal decoding accuracy is significantly higher than spatial decoding (F_(1)_=4.2, p=0.05).

### ‘Came From’ decoding weakens while ‘Go To’ decoding strengthens over the full 1-2-3 segment

To further investigate whether the transformation from ‘Came From’ to ‘Go To’ encoding occurred gradually or quickly, we examined the ‘Came From’ and ‘Go To’ decoding strength over the full 1-2-3 segment. We found that in the hippocampus, decoding varied across decoding categories (‘Came From’ vs. ‘Go To’) and segments (1-2 and 2-3; F_(1)_=8.9; p=0.006). Specifically, hippocampus ‘Came From’ decoding accuracy was significantly stronger than ‘Go To’ (F_(1)_=12.4; p=0.002), and the overall decoding was significantly stronger in segment 1-2 than 2-3 (F_(1)_=5.1; p=0.033), suggesting hippocampus was better at recalling memory of previously visited zones. Further, ‘Came From’ decoding was significantly stronger in segment 1-2 than in 2-3 (t_(21)_=-2.16, p=0.044; Fig. 3a *Top*), suggesting that the ‘Came From’ signal faded as the rats moved further away from zone 4 vs. 5. A similar dìerence in ‘Came From’ decoding accuracy across the 1-2-3 segment was not observed in PC (t_(28)_=-1.41, p=0.17; Fig. 3a *Bottom*) and there were not significant dìerences in ‘Go To’ decoding for hippocampus or PC across the 1-2-3 segment.

**Fig. 3:**
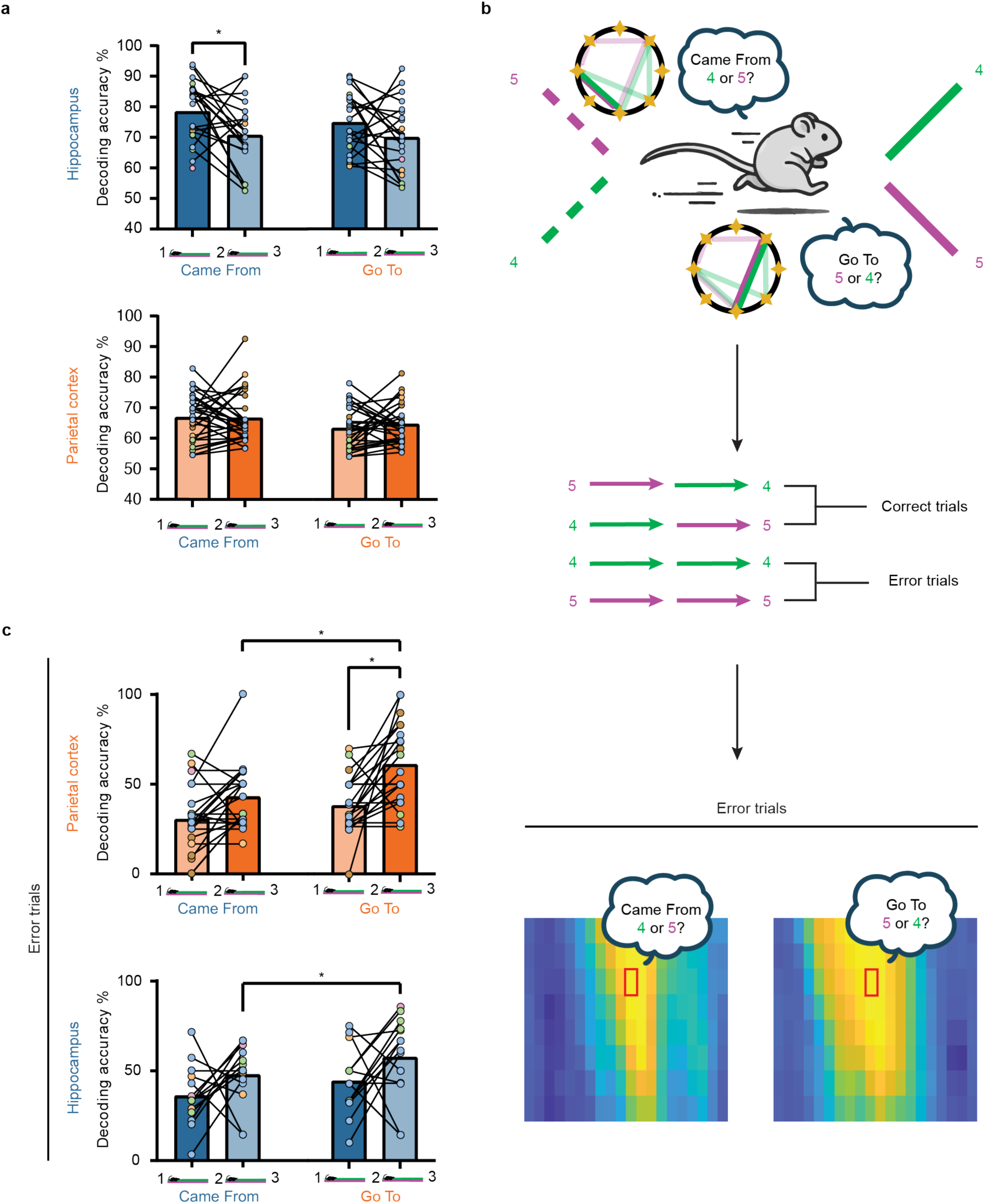
Over the Full 1-2-3 Segment, ‘Came From’ Decoding Becomes Less Accurate While ‘Go To’ Decoding Becomes More Accurate. **a**, Hippocampal (*top*) and PC (*bottom*) ‘Came From’ and ‘Go To’ decoding varies across segments 1-2 and 2-3, with hippocampal ‘Came From’ being less accurate during segment 2-3. **b**, Schematic illustrating the two ways in which error trials can be decoded. ‘Came From’ decoding can suggest that the rat may have remembered the wrong zone, predicting that it will make an error. In contrast, ‘Go To’ decoding can predict an error by indicating that the rat will ‘Go To’ the wrong zone. Note that the latter case could arise either because the rat remembered the wrong zone or selected the wrong action (i.e., ‘Go To’ decoding is better suited for predicting errors). **c**, Error trial prediction from decoding data for segments 1-2 and 2-3 varied significantly for PC MUA. Specifically, error decoding for ‘Go To’ was more accurate for segment 2-3, and error trial decoding was also more accurate for ‘Go To’ than for ‘Came From’ decoding in the 2-3 segment. Error trial decoding also varied significantly across the 1-2 and 2-3 segments for hippocampus, with error trial decoding being more accurate for ‘Go To’ than for ‘Came From’ decoding for the 2-3 segment. \**P* < 0.05; \*\*\**P* < 0.001.

Next, to further assess ‘Go To’ and ‘Came From’ decoding accuracy across the 1-2-3 segment, we evaluated the predictive accuracy of decoding for behavioral errors. Errors could result from “remembering” the wrong ‘Came From’ zone (i.e., ‘Came From’ decoding predicts a dìerent zone than the one the rat actually came from) or failing to transform that information into the correct future action (i.e., decoding predicts that the rat will ‘Go To’ the wrong future zone). Such patterns in the decoding data predict that the rat will make a future error (Fig. 3b). Note, because ‘Go To’ decoding can capture both errors, it should be more accurate at predicting errors than Came From’ decoding. Consistent with this expectation, PC error decoding varied significantly across decoding types (‘Came From’ vs. ‘Go To’) and segments (1-2 vs. 2-3; F_(1)_=5.0; p=0.034; p<0.001) and specifically for segment 2-3, ‘Go To’ was significantly stronger than ‘Came From’ (hippocampus: t_(14)_=3.40, p=0.009; PC: t_(22)_=2.48, p<0.001; Fig. 3c). In addition, we found that the ‘Go To’ error decoding was significantly better in segment 2-3 than in 1-2 for PC (t_(22)_=3.20, p=0.004; Fig. 3c). However, hippocampus error decoding did not vary significantly across decoding types (F_(1)_=0.43, p=0.51). While in PC, but not hippocampus, the overall decoding accuracy is significantly higher in segment 2-3 than in 1-2 (PC: F_(1)_=17.0, p<0.001; hippocampus: F_(1)_=3.6, p=0.078). This suggests that the ‘Go To’ signal in PC gets stronger over the full 1-2-3 segment. Overall, these findings provide evidence that the transformation from ‘Came From’ to ‘Go To’ is a gradual process, with fading of the ‘Came From’ signal and strengthening of the ‘Go To’ signal over the full 1-2-3 segment.

### ‘Came From’ decoding may emerge from PC route-centered encoding when the PC signal appears before the signal in hippocampus

Next, we sought to understand why in segment 1-2 sometimes hippocampus was leading, while at other times PC was leading. Previously, Nitz^25^ showed that PC single units map space in a novel ‘route-centered’ framework, meaning position information along a route. In the complex sequence task, during segment 1-2-3, the path is overlapping when the rat is coming from zone 4 or from zone 5, but it is possible that PC is splitting the traversal into separate routes (i.e., ‘route-centered’ encoding), similar to PC dìerentiation of loops on a spiral maze^26^. So, we plotted spike firing rate after z-score normalization for hippocampal (Fig. 4a *bottom*) and PC single cells (Extended Fig. 3a) and found that single cells were dìerentially active during the 1-2 segment based on when the rat had come from zone 4 vs. zone 5. Next, we generated similar plots for PC MUA (Fig. 4a *Top*) and found that in PC MUA, firing rate also dìerentiated ‘Came From’ zone 4 vs. zone 5 during the 1-2 segment (Fig. 4a *Top*). Consistent with the observations for single cells and MUA we found that the population of PC MUA clusters are less similar in segment 1-2 than 2-3 both when PC leads (t_(345)_=-2.77, p=0.05, Fig. 4b *Top Left*) and when hippocampus leads (t_(345)_=-2.29, p=0.002; Fig. 4b *Top Right*). Thus, it is possible that when hippocampus leads, such route-like encoding in PC was enabled by place cells in the hippocampus. A similar route-like encoding with dìerentiation during the 1-2 segment was not observed in the hippocampus (Fig. 4a & 4b *Bottom*). Instead, consistent with previous reports hippocampus encodes specific spatial locations (Fig. 4a *Bottom*).

**Fig. 4:**
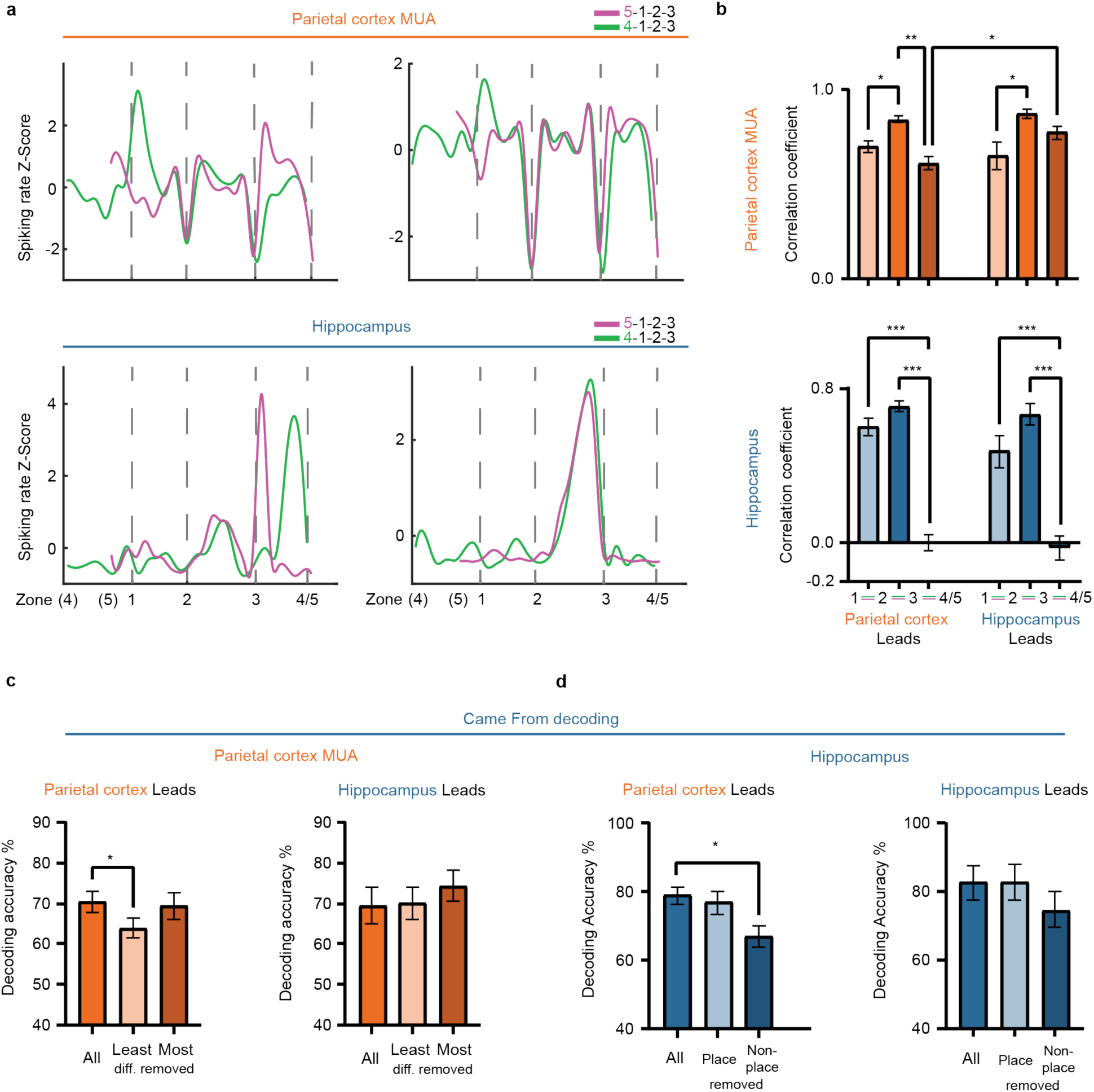
Route-centered Coding in PC When PC Leads and Place Cells in Hippocampus when Hippocampus Leads May Underlie ‘Came From’ Decoding. **a**, Examples of activity rate z-score normalized to mean firing rate and split by path 5-1-2-3 (magenta) and 4-1-2-3 (green) from PC MUA (*top*) and hippocampal single cells (*bottom*). Many PC MUA clusters dìerentiate the two routes during the 1-2 segment, while hippocampal single units tend to fire at a particular place on the maze, with the population of units covering the space of the maze. **b**, Correlation of PC MUA activity rate between routes 5-1-2-3 and 4-1-2-3 for each segment (1-2, 2-3, etc.) separated by behavior session type: PC leads vs. hippocampus leads for ‘Came From’ decoding during segment 1-2 (*top*) and hippocampus (*bottom*). PC, but not hippocampus, is more dissimilar for the 5-1-2-3 vs. 4-1-2-3 routes during segment 1-2 than 2-3. Note: 3-4/5 involves traversing physically dìerent locations, and both PC and especially hippocampus are also dissimilar for the two routes for this segment. **c**, ‘Came From’ decoding with PC MUA in behavior sessions where hippocampus (*left*) or PC (*right*) leads for ‘Came From’ decoding during segment 1-2. Removing route encoding PC MUA impairs decoding, but only when PC is leading hippocampus for ‘Came From’ decoding during segment 1-2. **d**, ‘Came From’ decoding with hippocampal single units in behavior sessions where the decoding peak in hippocampus (*left*) or PC (*right*) leads in segment 1-2. Removing non-place cells impairs decoding but only when PC is leading, suggesting that PC conveys ‘Came From’ information to non-place cells in hippocampus. Interestingly, removing place or non-place cells does not impair decoding when hippocampus is leading, suggesting both populations contain ‘Came From’ information, possibly because hippocampal place cells initially encode ‘Came From’ and then relay this information to non-place cells (as when PC is leading). *Error bar,* mean ± SEM; \**P* < 0.05; \*\*\**P* < 0.001.

To test if the route-like encoding we observed in PC MUA might underlie the ‘Came From’ decoding when PC was leading hippocampus, we classified the multiunit clusters from all data sets (133 clusters from rats 2-5) and then extracted the highest two-thirds correlation coefficients for segment 1-2 (i.e., least route dìerentiating clusters) and the lowest third (i.e., most dìerentiating clusters). We found that for behavior sessions where the PC decoding peak for ‘Came From’ during the 1-2 segment occurred before the hippocampus peak, removing the most dìerentiating clusters (t_(7)_=3.64, p=0.008), but not the least dìerentiating clusters (t_(8)_=0.20, p=0.85), significantly reduced decoding accuracy (Fig. 4c *Left*). However, removing the most dìerentiating clusters did not alter decoding accuracy when the hippocampus peak for ‘Came From’ decoding was before the PC peak (t_(6)_=0.72, p=0.50; Fig. 4c *Right*). Because we pooled across data sets to generate a list of correlation values for the 1-2 segment, sometimes within a data set one category contained more MUA clusters than another. Therefore, as an additional control to ensure that the impaired decoding was not simply due to removing more clusters in these instances, we also randomly selected and removed the same number of clusters as there were for the most dìerent 1/3 for each data set and decoding was not significantly altered (Extended Fig. 3b). Together, this data suggests that the route dìerentiating PC clusters contained information about the zone the rat came from, and that ‘Came From’ decoding when PC was leading hippocampus was driven by this route information originating in PC.

### ‘Came From’ decoding in PC may be relayed to hippocampal non-place cells when PC is leading, whereas when hippocampus is leading, both place and non-place cells in hippocampus contain sufficient information to accurately decode the ‘Came From’ location

Finally, to understand what might underlie ‘Came From’ decoding when hippocampus was leading PC, we classified hippocampal place cells (i.e., cells with significant spatial information and coherence^27^) and assessed their impact on ‘Came From’ decoding. We found that 72 out of 207 hippocampal single cells met the criteria for place cells. Removing non-place cells significantly impaired ‘Came From’ decoding in hippocampus when PC was leading (t_(9)_=-3.69, p=0.005; Fig. 4d *Left*), while removing place cells did not significantly alter decoding accuracy (t_(9)_=1.11, p=0.30), suggesting that PC route encoding MUA relays ‘Came From’ information to hippocampal non-place cells. To test if this impairment was simply caused by removing more cells (since there are more non-place cells), we randomly selected the same number of cells and found that decoding was not impaired (t_(9)_=-1.25, p=0.24; Extended Fig. 3b). However, when hippocampus was leading, removing either place cells or non-place cells did not impair ‘Came From’ decoding (t_(7)_s<1.38, ps>0.21; Fig. 4d *Right*), suggesting that in this case ‘Came From’ information was present in both populations of cells and that having only one source of information was sùicient for accurate decoding. These findings suggest that PC is feeding route-centered ‘Came From’ information into hippocampal non-place cells. When hippocampus is leading, ‘Came From’ information is present in both place and non-place cells. Together, this pattern of results could mean that in both cases, ‘Came From’ information is fed to non-place cells in hippocampus, when hippocampus is leading place cells are the source of this information and when PC is leading route encoding MUA is the source of this information.

### Egocentric tuning to future goal locations may underlie the ‘Go To’ decoding signal observed first in PC and then in hippocampus

PC and then hippocampus exhibited high prediction accuracies for ‘Go To’ decoding during the 2-3 segment (Fig. 2b, d), so we next sought the source of this encoding of future goal location (zone 4 vs. 5). We previously showed that PC single cells encode egocentric direction of goal locations^10^. Meanwhile, a number of groups have recently found egocentric encoding in hippocampal single cells, including egocentric encoding of goal locations^17–19^. Thus, we assessed egocentric tuning to the future goal location (zone 4 vs. 5) during the 2-3 segment.

As in Wilber et al. for PC^10^ and Ormond and O’Keefe for hippocampus^17^, we defined the egocentric direction of the future goal location as the angle between the vector of the rat’s head direction and the vector from the rat to the goal direction (Fig. 5a). Occupancy was then computed in segment 2-3 (Extended Fig. 4a), showing that most but not all of the egocentric positions of the goal location fell into the ranges of relative head directions that would occur if the rat followed the straight line between zone 2 and 3 with its nose always pointed towards zone 3. Firing rate was then computed for the egocentric direction of the goal location for zone 4 and zone 5 while the rat traversed the 2-3 segment. Hippocampal single cells, and PC MUA and single cells that demonstrated significant tuning to a specific egocentric goal location were identified (Fig. 5b; also see Methods, Extended Fig. 4b, c). For hippocampus, out of 180 cells, most cells had significant egocentric tuning with most tuned only to zone 4, and some tuned only to zone 5, and others were tuning bilaterally for both going to zone 4 and 5. Out of 162 PC MUA clusters, more than half had egocentric tuning. While a large number of cells this is in contrast to modular motion state encoding where nearly every module was turned to the motion state of the rat^23^. For PC MUA with egocentric tuning to future goal locations, most were tuned bilaterally for either zone 4 or 5, some were tuned only to zone 5, and a few were only tuned for going to zone 4 (Extended Fig. 5a). Similarly, we found that a notable proportion of hippocampal cells (46%) and PC MUA clusters (46%) exhibited distance tuning, although this was less prevalent than egocentric tuning. While distance tuning has been previously reported for hippocampal single cells, this is the first instance of it being observed in PC MUA (Extended Fig. 6). Next, we assessed the populations of egocentric tuning in hippocampal single cells and PC MUA clusters and found that the mean directions of both populations reflected the current future goal location, going to 4 or going to 5 (Fig. 5c). Furthermore, PC ‘Go To’ decoding was significantly reduced after removing egocentrically tuned PC MUA clusters (t_(12)_=5.39, p<0.001; Fig. 5d *Left*) and hippocampus decoding was impaired when egocentrically tuned hippocampal single cells were removed (t_(13)_=3.31, p=0.006; Fig. 5d *Right*). Neither decoding was impaired when non-egocentrically tuned cells or clusters were removed (PC: t_(12)_=0.89, p=0.39; hippocampus: t_(13)_=-1.44, p=0.17;). Furthermore, decoding was not impaired by randomly removing same number of PC MUA clusters (t_(12)_=0.89, p=0.39; Extended Fig. 4d *Left*) or hippocampal single cells (t_(13)_=-1.44, p=0.17; Extended Fig. 4d *Right*). Thus, egocentric tuning to future goal locations in PC may underlie ‘Go To’ decoding and that this information is then relayed to single cells in hippocampal single which also underlie ‘Go To’ decoding that is subsequently apparent in hippocampus.

**Fig. 5.**
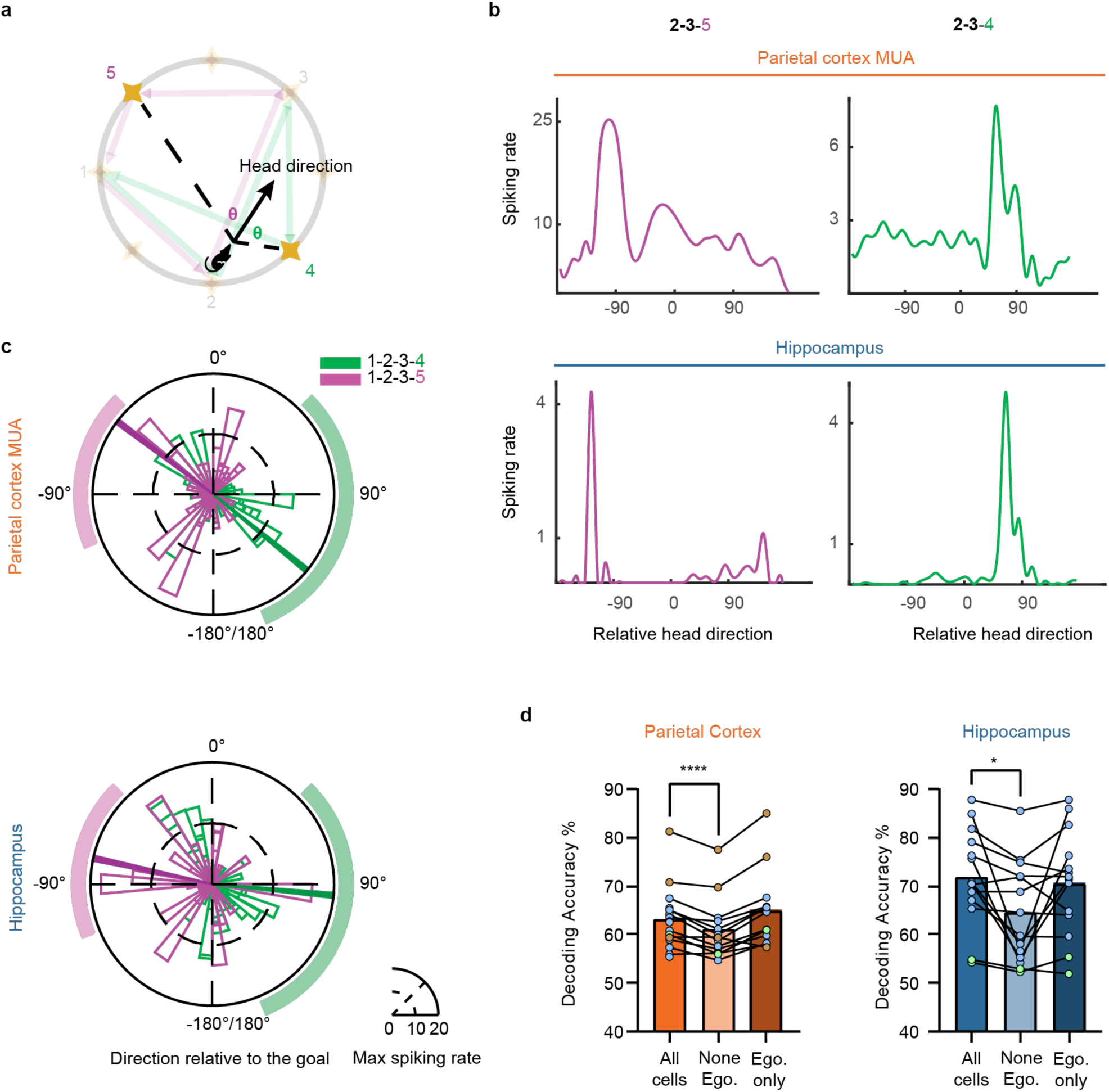
Egocentric Tuning to Future Goal Locations During the 2-3 Segment Underlies ‘Go To’ Decoding in PC and Subsequently in Hippocampus. **a**, Schematic of relative goal location fixed with respect to the current head direction (i.e., egocentric goal direction). **b**, Examples of egocentric tuning in PC MUA (*top*) and hippocampal single cells (*bottom*) split by path 1-2-3-4 (*left*, green) and 1-2-3-5 (*right*, magenta). **c**, Polar plots showing the egocentric tuning directions (peak) of each tuned cell or MUA cluster for PC (*top*) and hippocampus (*bottom*), divided by the path the rat will take 1-2-3-4 (green) and 1-2-3-5 (magenta). Shaded curve bars represent the ranges of relative head directions if the rat followed a straight line between zone 2 and 3, with its head is always orientated towards zone 3 during traversal. Occupancy data suggests that this is often, but not always the case, as expected since rats’ behavior is not stereotyped to this extent (Extended Figure 3a). Solid thick lines denote the mean directions of the egocentrically tuned population. **d**, ‘Go To’ decoding accuracies of PC (*top*) and hippocampus (*bottom*) varies when egocentrically tuned (or not tuned) cells or MUA are removed. Specifically, there is a significant reduction in decoding accuracy when egocentrically tuned cells or clusters are removed, but not when non-egocentrically tuned cells or clusters are removed. \**P* < 0.05; \*\*\**P* < 0.001.

All results up to this point are from the non-cued (memory) trials (see Methods); however, we also conducted separate analyses for cued trials. Until this point, no dìerences were observed for cued trials (Extended Fig. 7), suggesting that rats may ignore the cue light and perform cued trials from memory. However, we did observe a dìerence in egocentric direction encoding for PC during the interleaved light cued trials, reported here. We found that fewer PC MUA clusters were tuned to egocentric goal locations during cued sequences than non-cued sequences (χ^2^=35.75, p<0.001; Extended Fig. 5a), but not hippocampal single units (χ^2^=6.44, p=0.09), suggesting that with the aid of light cues, rats were less dependent on egocentric encoding of future goal locations. However, no reduction in PC (or hippocampus) future goal location distance encoding was observed for cued sequences. Further, there was no change in decoding of future goal locations (PC: t_(28)_=0.35, p=0.72; hippocampus: t_(21)_=1.14, p=0.27). Together, this suggests that ‘Came From’ encoding was unaltered by cue lights while ‘Go To’ egocentric encoding was reduced, but not to the point that it impacted decoding of the future goal location.

### Hippocampus and PC encode in their known allocentric and egocentric reference frames

Given the unique encoding we observed on this task, particularly for PC, we wondered if PC and hippocampus were behaving dìerently on this task or if traditional encoding states were still present. To start, hippocampal place cells were present, suggesting traditional encoding in this allocentric reference frame in hippocampus. Two examples of such place cells with activity in segments 3-4 or 3-5 are shown (Extended Fig. 8a). Thus, we used a leave-one-out decoding approach to decode full trajectories through the 1-2-3-4 and 1-2-3-5 segments and found that hippocampal single cells were able to predict trajectory accurately, including accurately predicting error trials (Extended Fig. 8b). Furthermore, when comparing the prediction performance of the whole sequence trajectory, hippocampal single cells outperformed PC MUA clusters (Extended Fig. 8c; path 1-2-3-4 X component of the path: t_(160)_=2.77, p=0.013; path 1-2-3-4 Y component of the path: t_(180)_=11.47, p<0.0001; path 1-2-3-5 X: t_(148)_=8.319, p<0.0001; path 1-2-3-5 Y: t_(148)_=8.549, p<0.0001).

Finally, to see if PC MUA activity patterns are consistent with our previous work^23^, which found that nearly all PC MUA clusters encode self-motion information, we found that PC MUA clusters encoded self-motion during the complex spatial sequence task. Two examples for a left and right turn tuned module are shown (Extended Fig. 8d). Similarly, we performed self-motion decoding by building a model from modular activity patterns from one of the two sessions and testing on the other session as in our previous work^23^. We found that PC could accurately decode the current self-motion state and that PC outperformed hippocampal single cells for decoding the current self-motion state (Extended Fig. 8e), a result that was consistent across behavior sessions (t_(18)_=7.12, p<0.0001; Extended Fig. 8f). Thus, in addition to the novel encoding states reported here, PC and hippocampus continue to encode allocentric and egocentric information respectively, consistent with our classical understanding of these brain regions.

## Discussion

In this study, we investigated the coordination between allocentric, route-centered and egocentric encoding in the rat brain by focusing on the hippocampus and PC, two “ends” of a larger brain network. Our findings reveal a surprisingly complex, coordinated, bidirectional flow of information between these regions during a relatively simple spatial sequence task, where rats had to remember a ‘Came From’ spatial location to select the appropriate ‘Go To’ spatial location (1-2-3-4-1-2-3-5). Firstly, both hippocampus and PC actively encode spatial memory and future action information, but this information is present in dìerent brain regions at dìerent times. During the 1-2 segment, hippocampal neurons sometimes encoded the spatial location that the rat had come from, which was subsequently evident in the PC MUA, and other times the signal emerged in PC before hippocampus, suggesting a flexible interaction between these regions. Examination of the encoding states underlying the spatial memory decoding (‘Came From’ zone 4 vs. 5) suggests two neural strategies are at play. When PC is leading, a ‘route-centered’ reference frame may underly ‘Came From’ encoding, which is then conveyed to non-place cells in hippocampus. Conversely, when hippocampus is leading, place-like encoding may underlie the decoding of where the rat had come from, which is then conveyed to non-place cells and PC. As the rats progressed to the 2-3 segment, the hippocampal ‘Came From’ signal fades, and the PC signal for the egocentric future goal direction is strengthened, with this egocentric signal subsequently detected in the hippocampus. In both PC and subsequently in hippocampus, this ‘Go To’ decoding signal is driven by encoding of the egocentric direction of the future goal location. This suggests that both neural strategies for encoding where the rat had come from converge on an egocentric signal in PC that is then relayed to hippocampus. In conclusion, our study shows that the hippocampus and PC form a dynamic network coordinating the relationship between previously visited spatial locations and egocentric plans to reach future goals. This process, which is marked by dynamic interactions between hippocampus and PC (and likely several brain structures in between), provides a window into the complex interactions and multiple reference frames that are likely to underlie navigation and spatial memory.

Navigation is a complicated process requiring coordinating information across multiple reference frames. Fluid navigation likely involves processing among numerous brain areas, selecting and shifting between many available strategies^28–30^. For example, many cell types are purported to underlie navigating our surroundings to achieve the current goal, and the density of these cell types varies across dìerent brain area. Some of these cell types encode in allocentric coordinates, spatial representation where locations are encoded relative to external landmarks rather than the individual’s current position, such as place cells^7^, landmark direction cells^10,31,32^, border cells^33,34^, grid cells^35^, and head direction cells^36^. These cells form a comprehensive neural representation of space, allowing animals to navigate and remember spatial environments.

The PC contains a mixture of egocentric encoding (e.g., locations and objects encoded relative to the individual), route-centered encoding, and allocentric (world-centered) encoding (e.g., head direction cells, or cells encoding a specific combination of head direction and the egocentric direction of a landmark/goal). The mixture of allocentric and egocentric encoding in PC positions it to participate in transforming between egocentric, route-centered, and allocentric representations of space, and in taking the output of allocentric to egocentric transformations and connecting this information to the execution of movements^3,6,10,23^. In addition, hippocampus and lateral septum^37^ have been reported to not only store spatial information, but also be involved in allocentric to egocentric transformations^38^. However, the present results suggest that this coordination between reference frames may actually occur across a much larger brain network than realized with recordings solely in hippocampus or lateral septum. Thus, the coordination between ‘Came From’ and ‘Go To’ we observed across the PC-hippocampal network is consistent with allocentric-egocentric transformations across this brain network. Further, the switching we observed between PC leading and hippocampus for ‘Came From’ decoding could reflect a shift in neural representations of space, perhaps switching between route-centered and allocentric (place-cell based) maps of space.

Consistent with the idea that animals can select from a range of navigation strategies, Nitz discovered a novel encoding reference frame that is neither allocentric or egocentric^39^, known as ‘route-centered coding’. This was was first observed in the PC^25^ and later also in the retrosplenial cortex^40^. Specifically, neurons respond to specific route segments, and can disambiguate position on a complex route, such as a spiral maze with repeated loops through similar segments. This coding can also be integrated with spatial and contextual cues in the retrosplenial cortex, creating a neural map that combines route and allocentric information. This integration helps dìerentiate the same route placed in two dìerent spatial locations, aiding in route planning and memory recall^41^. Neuroimaging studies by Maguire et al.^42^ support the idea that similar encoding may support route navigation in humans, who have PC activation during route navigation tasks. Here, we also observed route-centered encoding in PC that disambiguated the 4-1-2-3 and the 5-1-2-3 routes during the 1-2 segment. This route-centered encoding seemed to underlie the ‘Came From’ signal in the PC, but only when this signal appeared in PC before hippocampus. Interestingly, this signal was relayed to the non-place cells in hippocampus, potentially suggesting integration with other information in hippocampus^38^. In fact, such integration may begin upstream of the hippocampus, consistent with combinations of route-centered encoding in conjunction with allocentric information observed in retrosplenial cortex^40^ and in our present study also in the PC.

Moreover, we found evidence for a new encoding scheme at the MUA level in PC: egocentric encoding of future goal locations. Note that the egocentric goal direction encoding in PC MUA reported here dìers from our previous reports of single cells tuned to a single specific egocentric direction of a landmark or goal^10^ because, at the MUA level, tuning is predominantly bilateral for future goal locations on the rat’s right or left. When combined with the route-centered encoding also observed at the MUA level, this suggests that encoding at the MUA level is much more diverse than appreciated in our previous work^23^. Similarly, the PC MUA encoding of a single future goal location (and population of PC MUA underlying decoding of future action) is consistent with Harvey et al.^43^, which found PC single cells predict future decisions on a virtual T-maze. Further, this egocentric future goal direction encoding seems to underlie the decoding of future ‘Go To’ action we observed that emerged in PC^10,23,44–47^ and subsequently appeared in hippocampus, suggesting that coordinating between ‘Came From’ and ‘Go To’ is more complex than simply transforming a previous spatial location in to the appropriate future egocentric action. Why this ‘Go To’ signal is conveyed back to hippocampus remains to some extent a mystery, but the flow of information from hippocampus to PC and PC to hippocampus in this task suggests this brain network does indeed operate as a brain network and not a unidirectional circuit.

The spatial sequence task we employed here is similar to the T-maze tasks used to discover splitter cells in the hippocampus. These cells, first identified by Wood et al.^48^, dìerentially fire based on the future direction or spatial context (where the animal had come from on the previous trial). Splitter cells are thought to provide insight into neural mechanisms for episodic memory formation and neural substrates of spatial navigation^49^. It is thought that there are at least two types of splitter cells: retrospective splitter cells that help animals remember where they have come from, and prospective splitter cells that dìerentiate where the rodent will go to. Ferbinteanu and Shapiro^50^ argue that splitter cells are crucial for encoding sequences of places, aiding in distinguishing dìerent routes or sequences of events. This context-dependent activity integrates spatial and temporal information, supporting coherent memory formation^51,52^. Additionally, splitter cells contribute to route planning and decision-making, influencing behavior and memory recall^53,54^. Finally, hippocampal lesions only impair performance on the T-maze when a delay of at least 2s is imposed, leading to context-dependent hippocampal activity during the delay period^54,55^. Our task imposes a similar delay with a two-segment common section. Consistent with splitter cell activity during the delay period, we found that ‘Came From’ decoding occurs near the segment starting point (when the animal has slowed to receive a reward). The two-segment common arm also allows us to spatially and temporally segregate retrospective and prospective encoding. Typically, studies of splitter cells have restricted analyses to cells that have a place fields^49^, but we included both place and non-place cells. These approaches allowed for identification of a temporal sequence of prospective encoding and revealed that this prospective code may originate in PC (or an “upstream” structure) and then conveyed to hippocampal non-place cells.

Though traditionally known for allocentric encoding (e.g., placed cells and landmark direction cells), recent reports have also found egocentric encoding in hippocampus^17–19, also see: 56^. Here, we used the same approach as described by Ormond and O’Keefe^17^ to assess hippocampal egocentric tuning of cells to future goal locations. We found egocentric tuning in 51% of all place cells and 54% of all hippocampal cells recorded (Extended Fig. 5a, b). This is lower than the 80% of place cells reported by Ormond and O’Keefe^17^, higher than the 32% of place cells and 19% of all hippocampal cells with egocentric tuning reported by Sarel et al.^18^, and similar to the 46% of all hippocampal cells by Jercog et al.^19^. This suggests that, as we observed here, strategy variation within the same task is likely. Strategy may also vary across tasks, and that these strategy dìerences may be reflected in variation in neural coding.

Our study is the first to examine ‘Came From’ decoding on the complex spatial sequence task. However, it is not the first to use the task which was developed by Bower et al.^22^ (hippocampus only recording) and later used by Euston and McNaughton^57^ (prefrontal cortex only recording). Surprisingly, the previous studies did not find evidence for ‘Go To’ encoding in prefrontal cortex or hippocampus. Our approach may explain the dìerence between our results and those of Bower et al. for hippocampal ‘Go To’ decoding. We used a decoding matrix with dìerent window sizes for both spatial and temporal information, while Bower et al. used a decoding model with fixed parameters on only spatial information, excluding data from the segment beginning and end. This likely obscured the pattern of data we observed, where spatial decoding frequently occurred near the beginning of the segment.

While not inconsistent with the present results, the lack of true ‘Go To’ encoding by prefrontal cortex single cells did suggest a potential confound that we addressed. Euston and McNaughton found that apparent ‘Go To’ encoding was explained by the path variation. However, this encoding was present at the end of the trajectory, not the beginning as in our data, and path variation was negatively correlated with PC decoding accuracy, but positively correlated with hippocampus decoding accuracy (Extended Fig. 9). Furthermore, we found that in segment 1-2, there were no significant correlations in either hippocampus or PC with path variation and ‘Came From’ decoding accuracy. Together, these results suggest that while interpreting hippocampal ‘Go To’ decoding with caution is warranted, path variation cannot account for the pattern of the present results.

We found evidence that the same type of information exists in hippocampus and PC but at dìerent times, suggesting coordination across a network involving the PC, hippocampus and several intermediate structures. We also found that the ‘Came From’ signal gradually faded and the ‘Go To’ signal gradually strengthened over the full 1-2-3 segments, again consistent with reference frame coordination being a network-level event. Thus, spatial processing in allocentric, egocentric and route-centered reference frames is not strictly divided but rather dynamically distributed across multiple brain regions. Furthermore, we provide evidence that neural strategy can rapidly switch within the same task, suggesting that rapid adaptation to the task’s demands and context is possible and potentially more frequent than appreciated in prior spatial navigation research. Our work thus òers new insights into the neural mechanisms of spatial navigation, highlighting the importance of considering multiple reference frames and navigation strategies across a brain network extending at least from PC to hippocampus to understand the full scope of spatial processing in the brain.

## Main Figures

**Extended Fig. 1:**
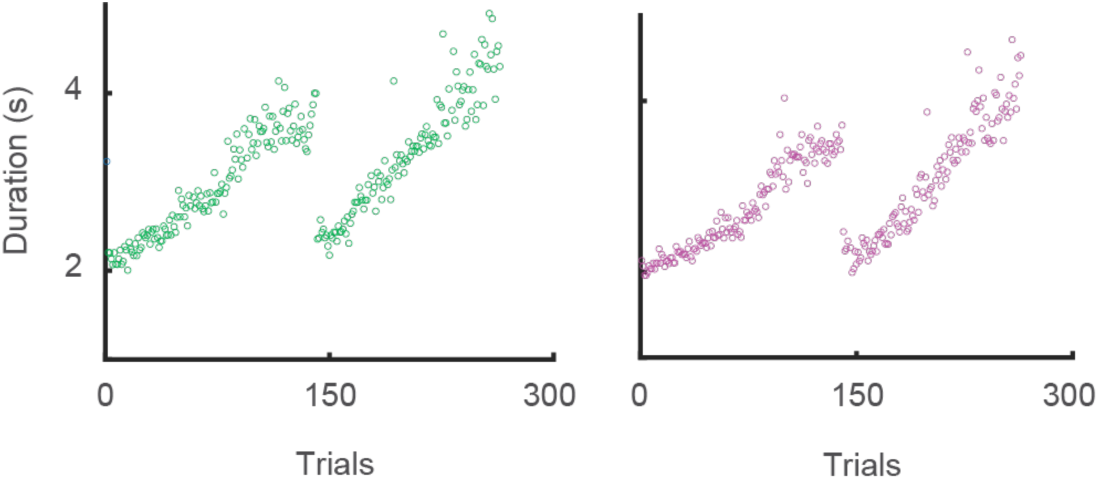
Running Durations Varied Within Sessions. Example of running durations in segment 2-3 split by path 1-2-3-4 (green, left) and 1-2-3-5 (magenta, right), showing that rats progressively run more slowly over the course of a recording session. To address this issue, we normalized time for temporal decoding.

**Extended Fig. 2:**
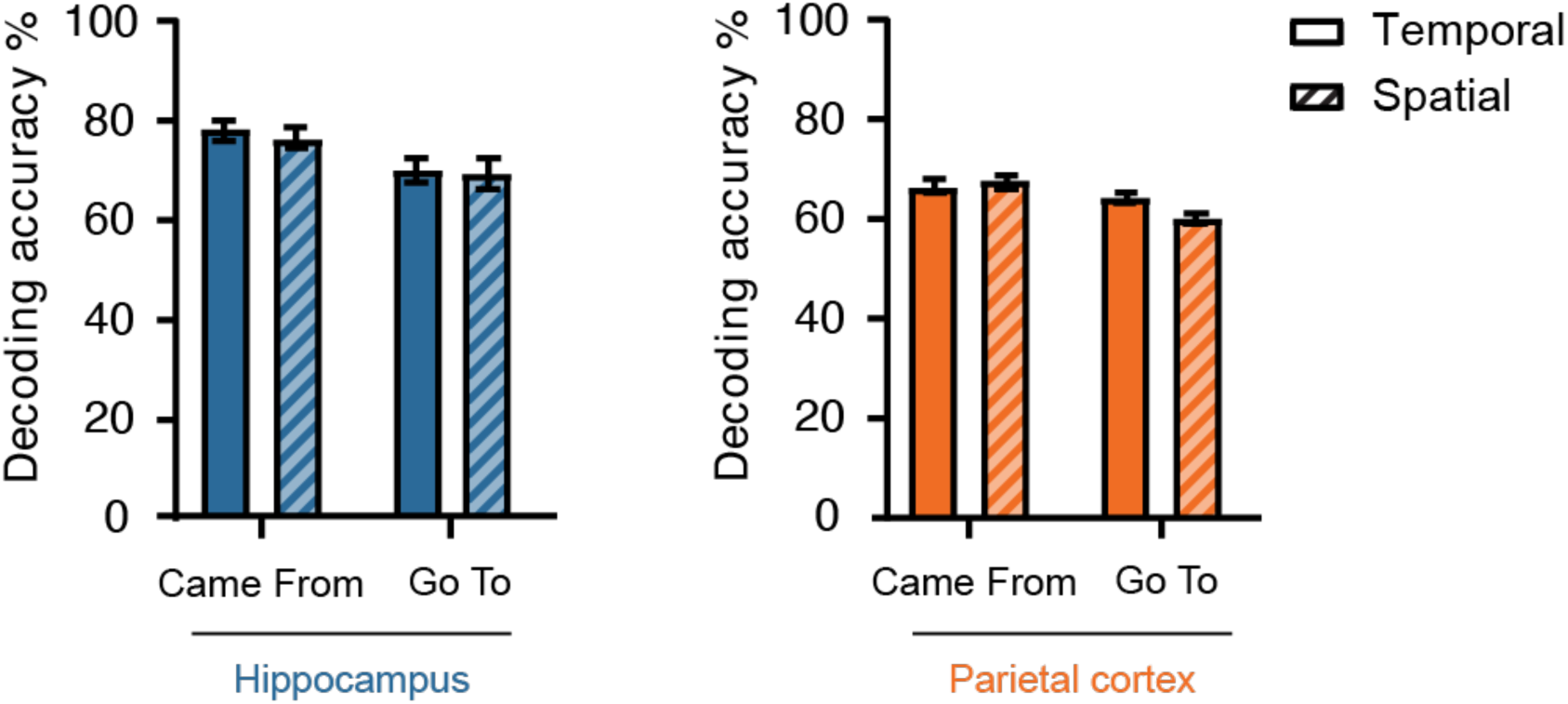
Decoding Accuracy is Generally Similar for Spatial and Temporal Decoding. Comparison of spatial and temporal decoding in hippocampus (*left*) and PC (*right*) for ‘Came From’ decoding and ‘Go To’ decoding. Temporal decoding accuracy is significantly higher than spatial decoding (F_(1)_=4.2, p=0.05). Additionally, for both hippocampus and PC, ‘Came From’ decoding is significantly stronger than ‘Go To’ (hippocampus: F_(1)_=4.4, p=0.05; PC: F_(1)_=5.7, p=0.02). *Error bars,* mean ± SEM.

**Extended Fig. 3:**
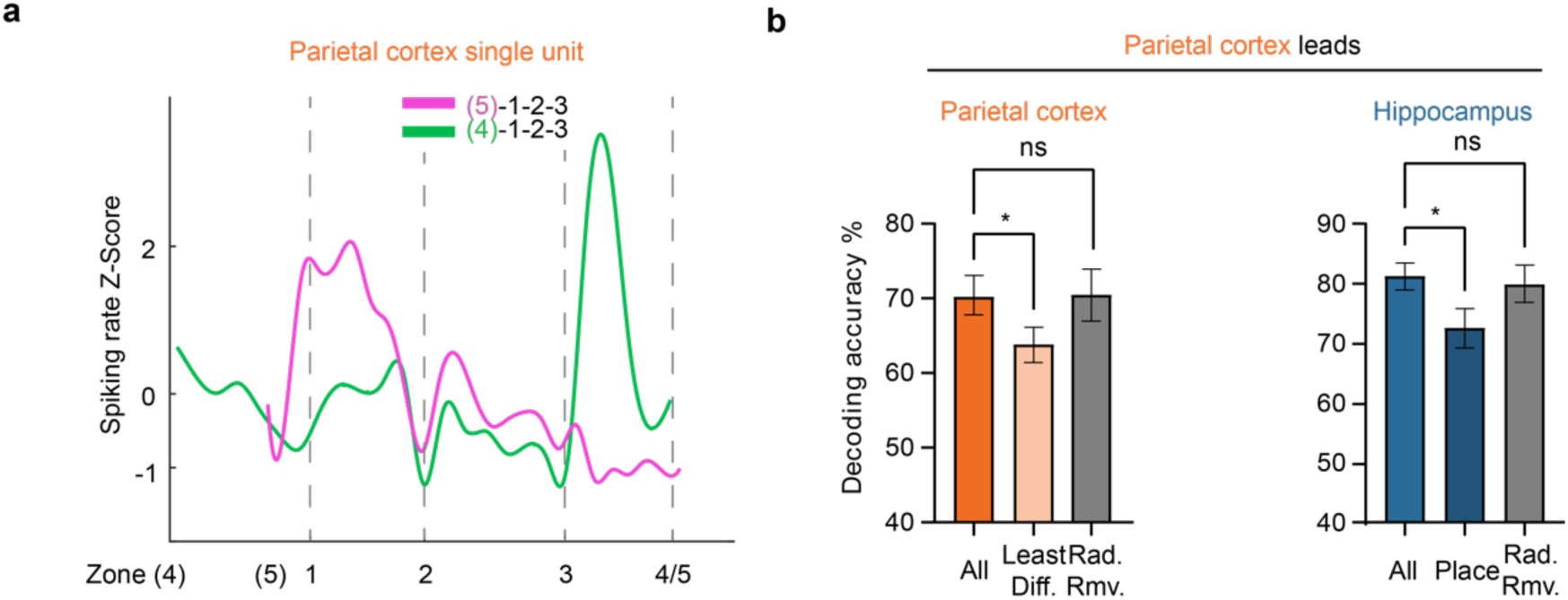
PC Single Units Also Encode the Route and Removing Equal Numbers of MUA Clusters or Single Cells Does Not Impair Decoding. **a**, Examples of PC single cell firing rate z-score normalized to mean firing rate, split by path (5)-1-2-3 (magenta) and (4)-1-2-3 (green), show that PC single cells also encode the route. **b**, ‘Came From’ decoding accuracies after removing the same number of PC MUA clusters or hippocampal single cells as were removed to produce the significant dìerence in Fig. 4c and d (replotted here), did not significantly alter decoding (PC: *P* = 0.76; hippocampus: *P* = 0.43). Least dìerentiating (Least Dì.). Randomly removed (Rad. Rmv.). *Error bars,* mean ± SEM; \**P* < 0.05.

**Extended Fig. 4:**
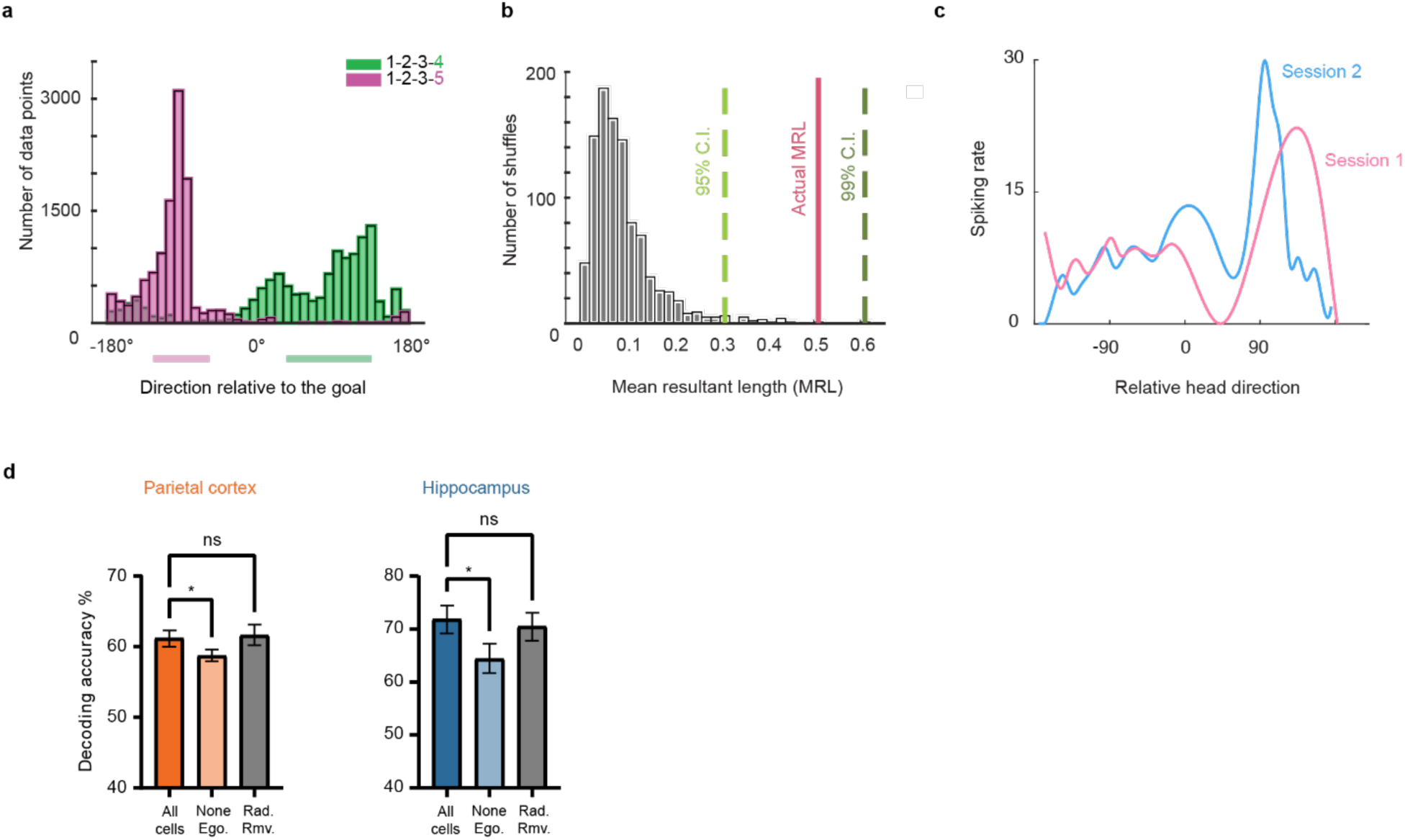
Criteria for Egocentrically Tuned MUA or Single Cells. Removing Equal Numbers of MUA Clusters or Single Cells Does Not Impair Decoding. **a**, Occupancy of relative head directions in one behavior session, split by path 1-2-3-4 (green) and 1-2-3-5 (magenta). Straight bars under the x-axis represents the range of relative head directions that would occur if the rat followed a straight line between zone 2 and 3 and its head was always pointed directly towards zone 3. **b**, Example of shùled distribution (1000 times of shùling position data) of mean resultant lengths for hippocampal single cells. **c**, Example of session stability test of PC tuning peaks (peaks between two sessions vary ≤ 40 ° as well as Rayleigh test p≤0.05). Note: Due to the issue that the shùle test for hippocampal single units was not able to handle high spiking frequencies of MUA (leading to false positives), we used the criteria for PC MUA that we have previously used for egocentric tuning in PC (significant Rayleigh test and stable directional tuning across sessions; see Methods^10^). **d**, ‘Go To’ decoding accuracies were not significantly reduced when removing the same number of egocentrically tuned PC MUA clusters or hippocampal single cells as in Fig. 5d (replotted here), compared to decoding with all clusters/cells (PC: *P* = 0.47; hippocampus: *P* = 0.17). *Error bars,* mean ± SEM; \**P* < 0.05.

**Extended Fig. 5:**
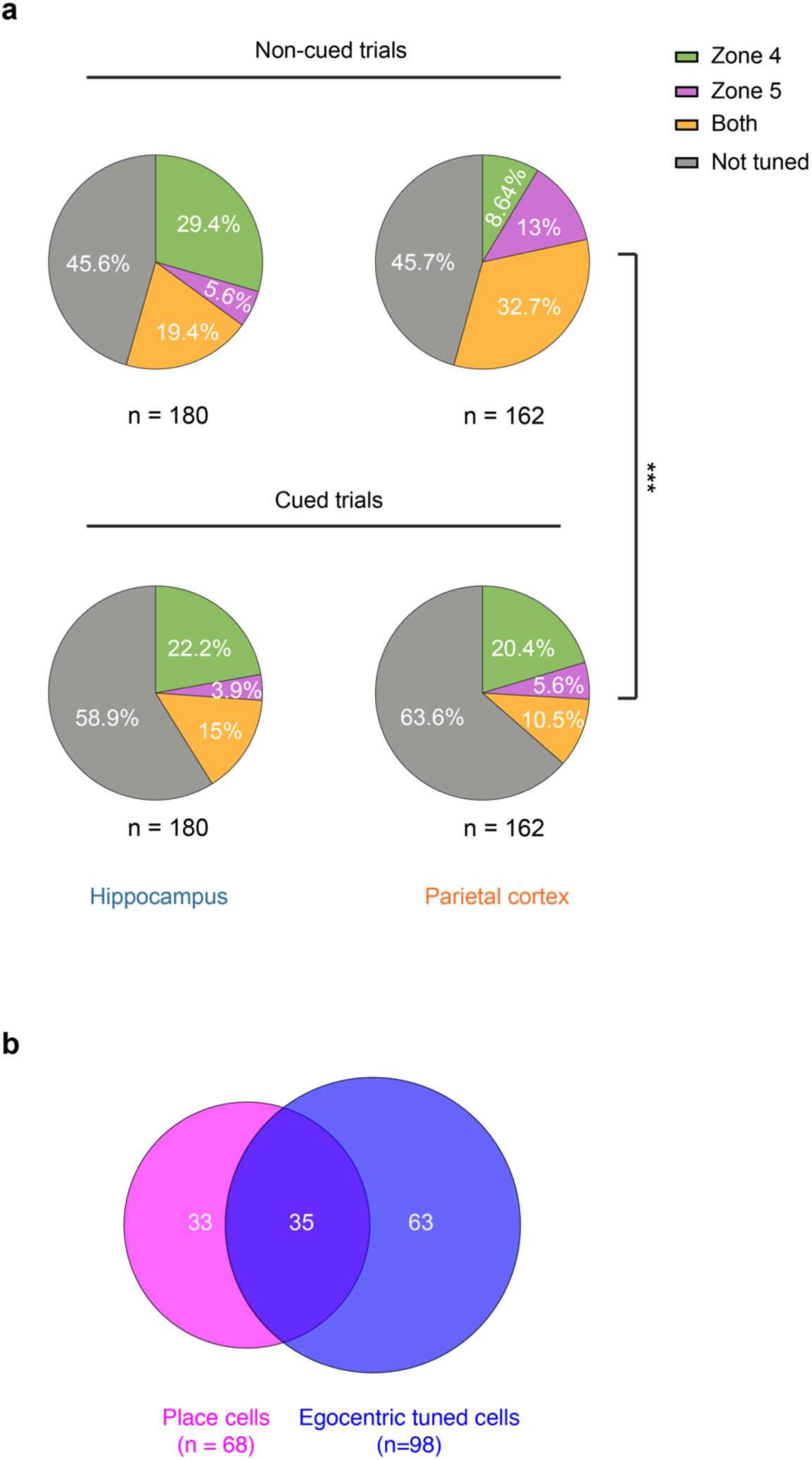
Proportion of Cells with Egocentric Tuning to Future Goal Locations. **a**, *Top*: Proportion of egocentric tuning type (Zone 4 only, Zone 5 only, Both Zone 4 and 5, or no egocentric tuning) during non-cued (memory) sequences for hippocampus and PC. *Bottom*: Same as *top*, but for light cued sequences (cue lights sequentially illuminate each zone in the sequence). **b**, Total numbers of recorded egocentric tuned cells and place cells during non-cued sequences in hippocampus. \*\*\**P* < 0.001.

**Extended Fig. 6:**
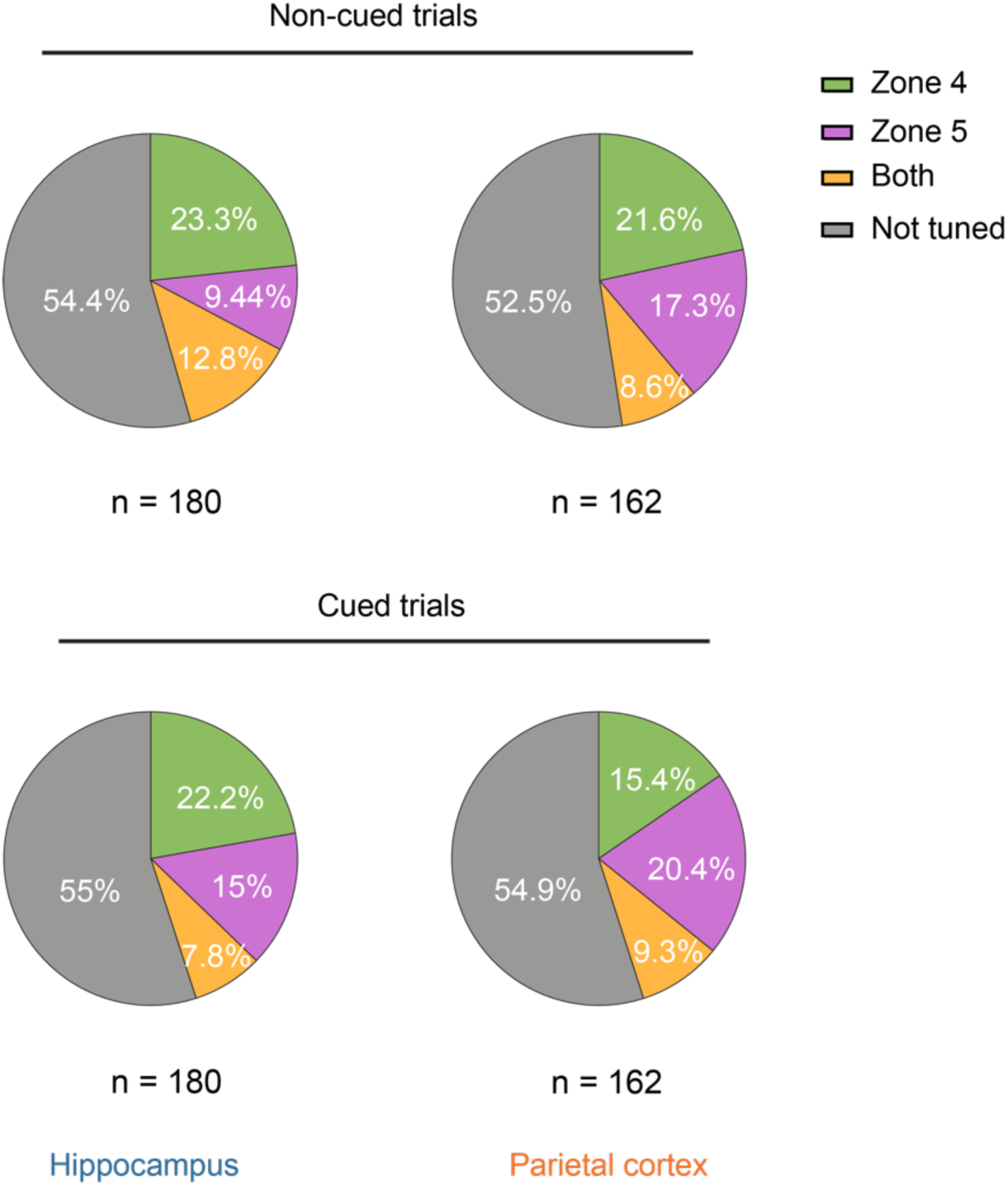
Future Goal Distance Tuning Proportion. *Top*: Proportions of distance-tuned cells for a single future goal location, or both future goal locations, during non-cued (memory) runs through the full sequence. *Bottom*: Same as *top*, but for light cued sequences.

**Extended Fig. 7:**
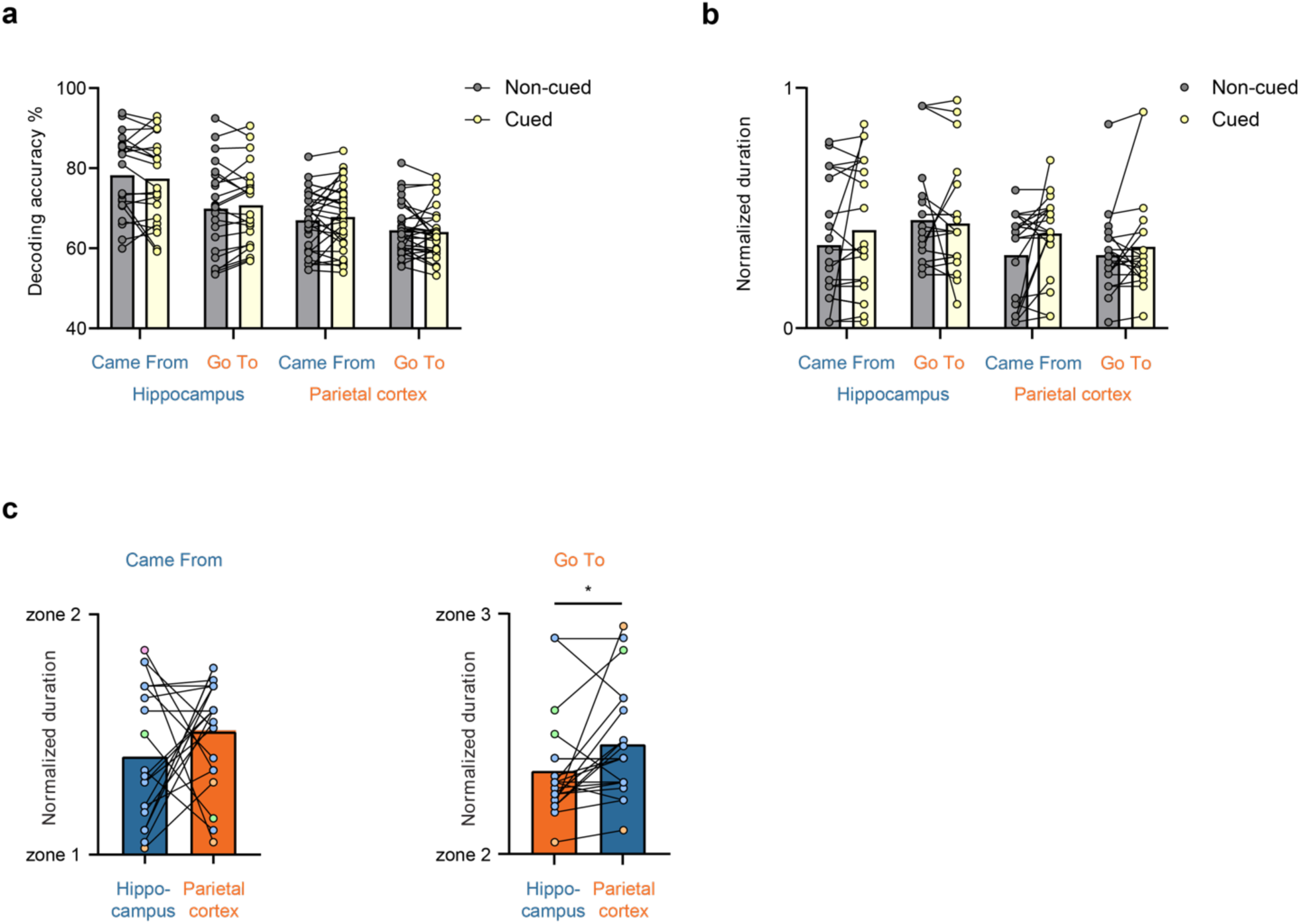
Temporal Decoding Results Were Not Different Between Cued and Non-Cued Trials. **a**, No dìerences between cued and non-cued trials for hippocampus and PC decoding accuracies for ‘Came From’ and ‘Go To’ decoding (hippocampus: t_(21)_s<1.14, *ps*>0.27; PC: t_(28)_s<0.37, *ps*>0.72). **b**, No dìerences in temporal relationships of decoding peaks of ‘Came From’ and ‘Go To’ from hippocampus and PC during cued and non-cued trials (hippocampus: t_(20)_s<1.54, *ps*>0.14; PC: t_(20)_s<1.49, *ps*>0.12). **c**, Temporal relationships for cued trials during segment 1-2 (*left*, t_(20)_s=-1.25, *p*=0.22) and 2-3 (*right*, t_(20)_s<-1.55, *p*=0.04) were consistent with non-cued trials (Fig. 2d).

**Extended Fig. 8:**
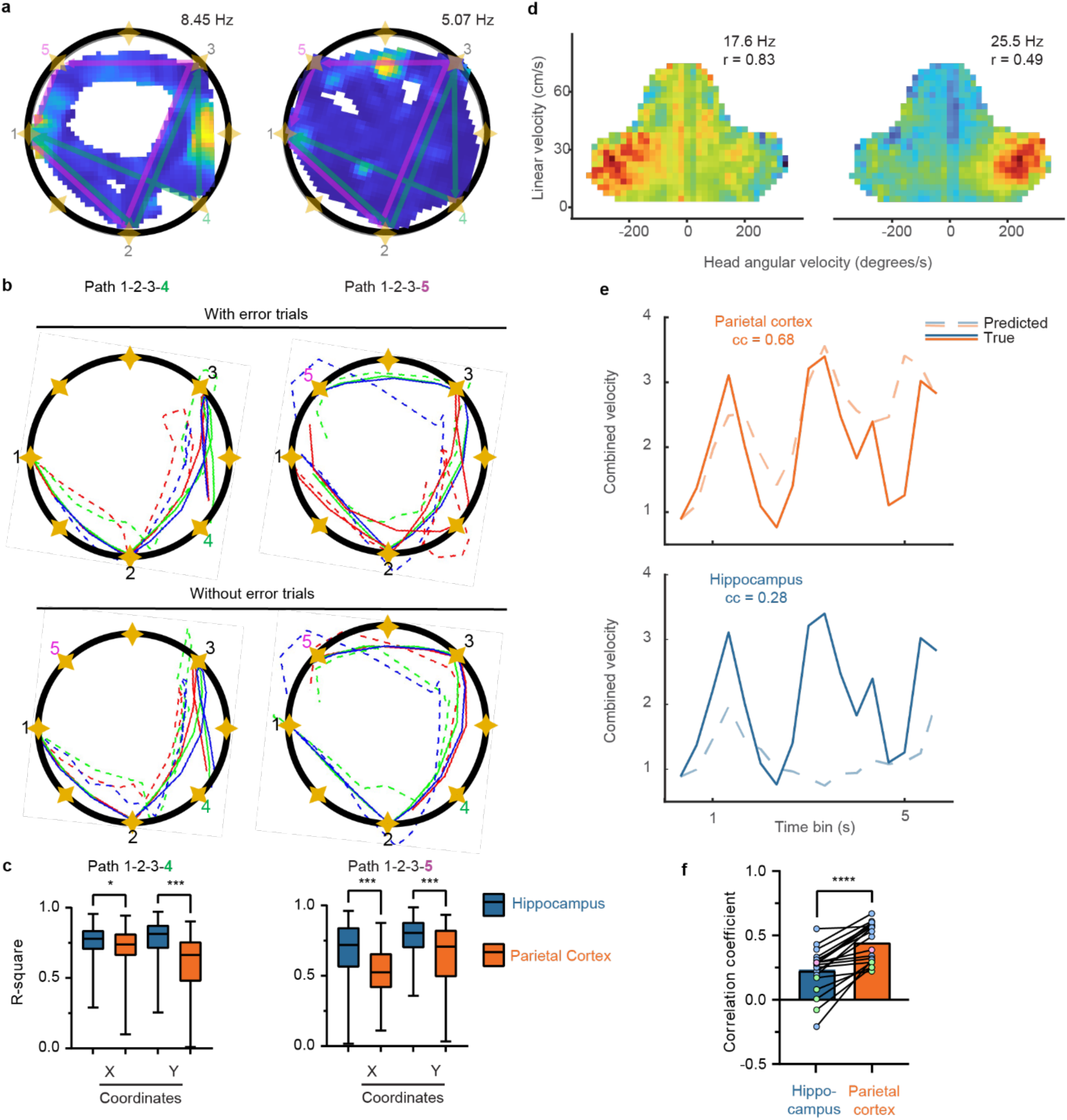
Allocentric Trajectories and Motion State Encoding Are Also Present in Hippocampus and PC. **a**, Examples of place cell firing maps with sequence paradigm overlaid, showing that some place cells dìerentiate the distinct places that are in the traversal from zone 3-4 and 3-5. **b**, A leave-one-trial-out approach was used to build a model for decoding whole sequence trajectories and was highly accurate both with (*top*) or without error trials (*bottom*). For illustrative purposes, only the trajectory decoding is shown separately for path 1-2-3-4 (*left*) and 1-2-3-5 (*right*). Solid line: true trajectory. Dashed line: predicted trajectory. **c**, R^2^ of correlation of predicted and true path plotted separately for the X and Y coordinates and split by path 1-2-3-4 (*left*) and path 1-2-3-5 (*right*). Whiskers: min to max. **d**, Examples of self-motion tuning PC MUA. **e**, Two-dimensional self-motion was transformed into one-dimensional overall motion (see Methods; as in our previous work^23^). A split-session decoding approach was used to predict the current self-motion state of the rat. Examples of self-motion prediction from PC MUA (*top*) and hippocampal single cells (*bottom*). Solid line: true motion. Dashed line: predicted motion. **f**, Correlation of predicted and true self motion is higher when the population of PC MUA is used than hippocampal single cells. \**P* < 0.05; \*\*\**P* < 0.001.

**Extended Fig. 9:**
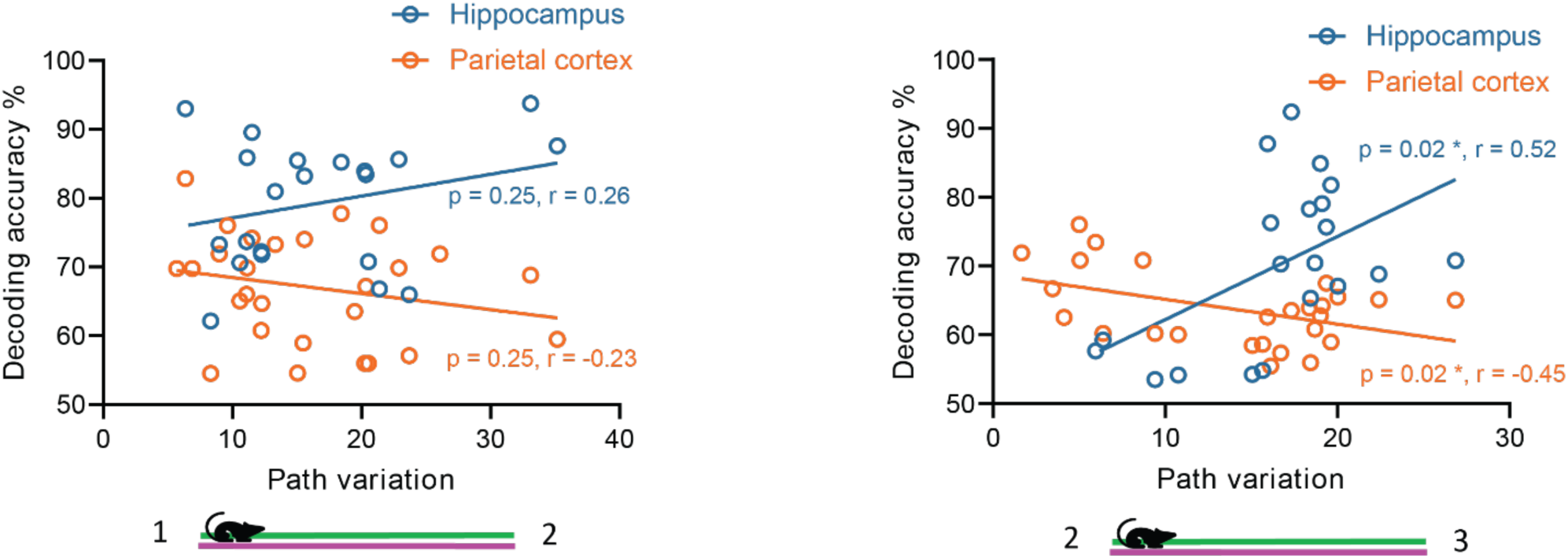
Path Variation Does Not Account for the Pattern of Results. Correlations between decoding accuracies and running path variation for hippocampus and PC for ‘Came From’ decoding during segment 1-2 (left) and ‘Go To’ decoding during segment 2-3 (right). Path variations were determined by the distance of individual paths from the mean path for the same segment for the two routes (going to 4 vs. 5). \**P* < 0.05

**Extended Fig. 10:**
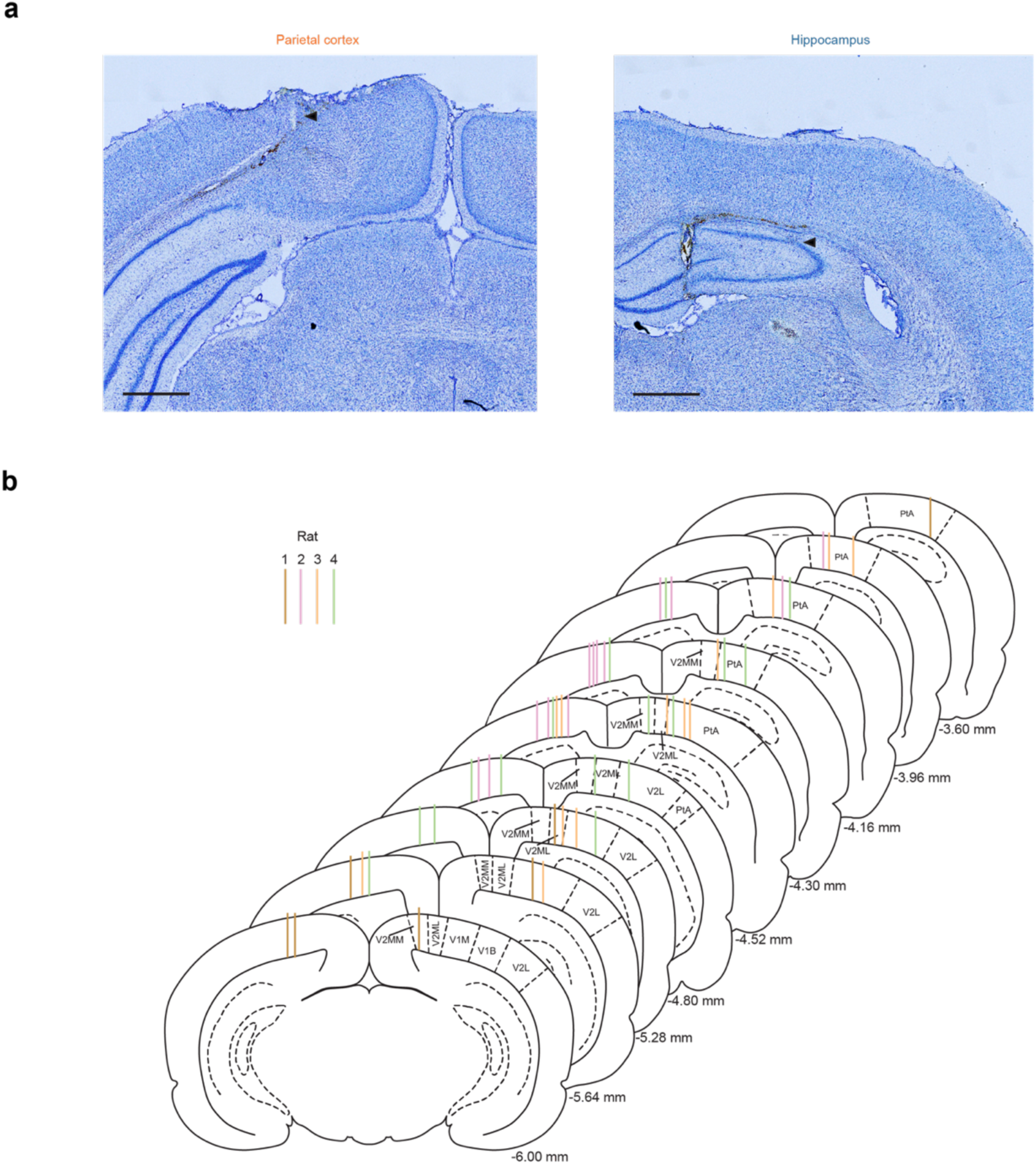
Recording Locations in Hippocampus and PC. **a**, Nissl-stained coronal section examples of the marking lesion for a tetrode positioned in PC (*left*) and hippocampus (*right*), with marking lesion location indicated by a black arrow. Scale bar, 1 mm. **b**, Coronal sections throughout the anterior (*top*) to posterior (*bottom*) extent of the rat PC^58^, color coded by rat 1-4. Rat 5 died unexpectedly, and thus reliable histology could not be performed. Tracts represent tetrode position with respect to the PC and hippocampus. Distance posterior to bregma is listed for each slice (lower right). Secondary visual cortex lateral area (V2L); medolateral area (V2ML); mediomedial area (V2MM); parietal association cortex (PtA); primary visual cortex (V1M); primary visual cortex binocular area (V1B). PtA, V2MM, and V2ML were considered parietal cortex based on our _prior work10,23,59,60._

**Extended Fig. 11:**
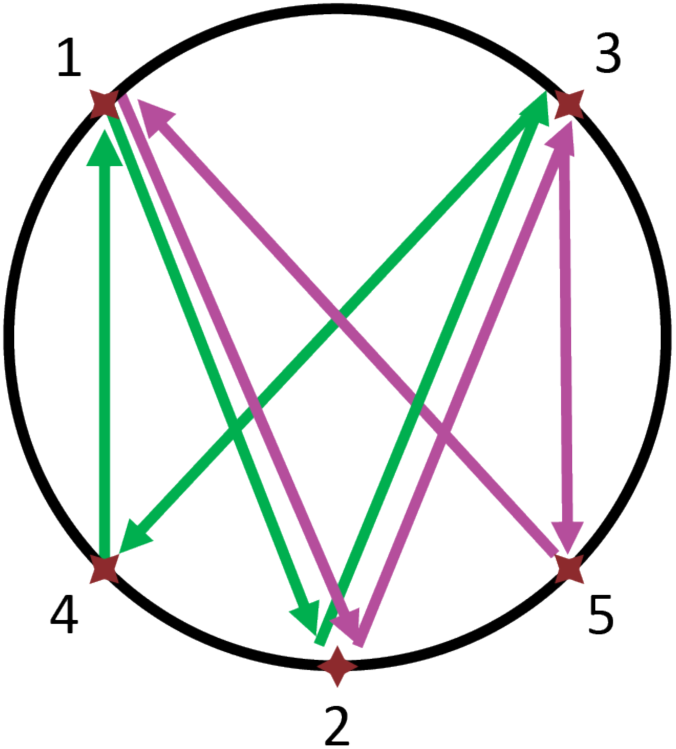
Sequence Schematic for Rat 2. In this variation of the spatial sequence that was used for rat 2, segments 1-2 and 2-3 are the same in length.

## Materials and Methods

### Subjects

Fischer/Brown Norway (F344BNF1) rats (n=5), comprising males (n=3) and females (n=2), aged 5-13 months, were housed in a 12:12 hour light/dark cycle. Rats were either food-deprived to 85% of their baseline weight and motivated with Ensure as a food reward (n=1), or received electrical stimulation of the medial forebrain bundle (MFB) as a reward (n=4)^10,22,23,57,61,62^. The rat receiving food reward underwent surgery for hyperdrive implantation, while the rats receiving brain stimulation reward underwent surgeries for both bilateral stimulating electrodes and subsequent hyperdrive implantation. For the latter, after one week of recovery, rats were trained to nose poke for MFB stimulation. Dìerent brain stimulation parameters were tested (200-500 ms, 100-300 Hz, 50-200 µA), and the optimal parameters that did not cause motor artifacts were selected for behavioral training and testing. Rats capable of performing more than 50 nose pokes per minute underwent a second surgery for hyperdrive implantation. Those unable to meet this criterion were discontinued from the study (n=8).

## Surgical Procedures

### MFB Stimulation

Bipolar stimulating electrodes were implanted bilaterally at the MFB (2.0 mm posterior from bregma, 1.7 mm from midline, 8.0 mm ventral from dura). MFB stimulation was necessary to obtain sùicient trials for some analyses. To ameliorate potential concerns about MFB effects on neural activity, data were excluded for the duration of the brain stimulation plus an additional post-stimulation 20 ms blackout period^10,22,57,61,62^. Further, in previous experiments where sùicient coverage for analyses was achieved using food reward (and consistent with the pattern of data in the present paper), identical results were obtained using both MFB stimulation and food reward, suggesting that the results obtained from MFB experiments are generalizable^57^.

### Recording Array

Custom 3D printed recording arrays^10,23,63^ were positioned over the bilateral PC (centered 4.5 mm posterior from bregma and 2.95 mm from midline)^10,23,59^. Each recording array contained 16-18 independently drivable tetrodes and 1-2 reference electrodes. Each tetrode consisted of four polyimide-coated, nichrome wires (14 µm diameter) twisted together. Beginning during the implantation surgery and continuing on subsequent days, each tetrodes targeting PC was turned down 31 µm and those targeting hippocampus were turned down 124 µm daily until they reached the hippocampal CA1 region based on both depth records and characteristic hippocampal local field potential (LFP). Then, each tetrode was turned every other day when appropriate to prevent getting stuck. Tetrodes targeting PC and hippocampus were distributed evenly across the left and right hemispheres (Extended Fig. 10).

### Recording Procedures and Data

Electrode interface boards (EIB-72-QC-Large) were used to connect the headstage (HS-72-QC-LED) to the recording system (Digital Lynx 16SX Neuralynx),^as in 24^. Each daily recording session included one 1-h rest session, followed by one 50-min behavior session, one 1-h rest session, another 50-min behavior session, and a final one 1-h rest session. Threshold spike waveforms were bandpass-filtered 0.6-6 kHz and digitized at 32 kHz. A continuous trace was simultaneously collected for processing as LFP from one of the tetrodes wires (bandpass-filtered 0.1-1000 Hz and digitized at 6400 Hz). Spike and LFP channels were referenced to an electrode in the corpus callosum. Rat position and head direction were tracked using red and blue LED lights on the headstage, and online position information was used to trigger automatic delivery of MFB stimulation or food rewards. Position and head direction data were collected at 30 Hz as non-interleaved video and co-registered with spikes, LFPs, and stimuli. At the end of recording sessions each day, tetrodes were turned, and configuration adjustments were made to allow stabilization overnight.

Spike data were automatically overclustered using KlustaKwik, then manually adjusted using a modified version of MClust (A.D. Redish). All spike waveforms with a shape, and across tetrode cluster signature suggesting that they were likely MUA and not noise, were manually selected and merged into a single MUA cluster (http://klustakwik.sourceforge.net^64^). For single-unit analyses only (not MUA analyses) three additional criteria were applied: 1) Only spike clusters with < 0.4% of the spikes in the first 2 ms on the autocorrelation were included. 2) Only spike clusters with < 15% threshold cut (as assessed by peaks plots) over the course of the entire 5 h recording session were included. 3) Only spike clusters with well-defined cluster boundaries were included for analyses.

Data recorded from a subset of the sessions were spike sorted and analyzed (rat 1 = 7 sessions, rat 2 = 1 sessions, rat 3 = 3 sessions, rat 4 = 4 sessions, and rat 5 = 14 sessions). All sessions containing both PC MUA and hippocampal single units from rat 2-4 were analyzed. Sessions were randomly chosen from a subset of all sessions containing PC MUA from rat 1 and hippocampal single units from rat 5. A subset was selected for rat 5 because this rat had so many high-quality data sets that analyzing all of them would have led to a larger imbalance in data sets per rat. A subset of the data sets was analyzed in rat 1 because we did not collect hippocampal single units from that rat. We analyzed data recorded in the dorsal CA1 field of the hippocampus simultaneously with the PC data (207 putative pyramidal cells recorded from the same 22 sessions as the PC data above for rats 2-5).

### Behavior

Very similar to the task used in our previous research^10^, training and testing took place on a large circular platform (1.5 m diameter) with 32 zones equally distributed along the platform edge, each with a light cue and a liquid Ensure feeder port. A custom program, which we have made freely available ^24^ (interfaced with the maze via a 48-channel USB-DIO-48 module through a Sysly IDC50-B breakout board), was used to control maze events and the delivery of either food rewards through feeders (valves are controlled using a 16-channel Digital I/O USB Module, USB-IDO-16) or MFB stimulation via a Stimulus Isolator (World Precision Instruments A365) when the rats entered a 10 cm diameter zone in front of the active/next zone in the sequence. The program also generated a coded timestamp in the Neuralynx system for each maze event.

Rats first performed the *alternation task*, where their movements were restricted by barriers to allow alternating between a pair of cue lights on opposite sides of the track. To ensure cues were visually salient, lights marking the reward zone were flashed at ∼3 Hz (with equal on/òtime) when that reward zone was activated (i.e., the current next zone in the sequence, in this case, the alternation sequence). The first light remained activated until the rat reached the reward zone and received MFB stimulation or food reward, after which the cue light in the opposite reward zone was activated.

If the rats were able to keep running back and forth to get rewards for 20 min, then they were advanced to the next *random lights task*. For this task, the whole platform was open (i.e., the barriers were removed), and one of the light cues positioned at one of the 32 zones was selected randomly and illuminated with the blinking cue light. The rat had to run to a spatial zone in front of the cue light to receive a reward. Once the rat ran to more than 10 zones/min for 30 min, it was advanced to the final task, the complex spatial sequence task. At this point, the recording session was also expanded to include three rest sessions (data not reported here) interspersed with two behavioral sessions on the apparatus, except for 2 of the 29 total sessions presented here, which had one behavioral session between two rest sessions.

### Complex Sequence Task

A spatial sequence was formed from 5 zones out of 32 and constructed with a repeated element: 1-2-3-4-1-2-3-5. Rats started from zone 1 to 2 to 3, then either went to zone 4 or zone 5. Two isomers of sequences were used. Rats 1 and 3-5 were run on a sequence where the lengths of segments 3-4 and 3-5 were identical (Fig. 1a). Rat 2 was run on a dìerent sequence of which the lengths of segment 3-4 and 3-5 were not identical, as is described in our and others previous work^22,23,61^(Extended Fig. 11). Rewards were delivered at each zone in the spatial sequence. Each behavioral session began with two runs through the 8-item sequence with cues. For these cued sequences, cue lights would lead the rat through the 8-item sequence one zone at a time. The two cued sequences were followed by two non-cued memory sequences, with no cue lights used unless the rat did not reach the rewarded zones before 18 seconds had elapsed. After 18 seconds, a cue light would illuminate the next zone, and the trial would be counted as an error to prevent rats from getting lost in the sequence. For rats 1 and 3-5, pairs of sequences were repeated, alternating between two cued sequences and two non-cued memory sequences until the 50 min behavior session time had elapsed. For rat 2, sets of three cued and three non-cued sequences were interleaved.

### Histology

After the final recording session, rats were deeply anesthetized with Euthasol and transcardially perfused with 1x phosphate bùered saline (PBS) followed by 4% paraformaldehyde in 1x PBS. The whole head was postfixed in 4% paraformaldehyde with electrodes in place for 24 h. Brains were then extracted and cryoprotected in 30% sucrose. Frozen sections were cut (40 μm) using a sliding microtome, mounted on positively charged slides, stained with cresyl violet, and imaged with a 10x objective (approximately 100x magnification) using a Zeiss Axio Imager.M2 microscope.

### Spatial Neural Decoding of ‘Came From’ Location and ‘Go To’ Location

A binary logistic regression was applied to decode the space-based signals. The spikes (MUA or putative single units) during zone 2 to 3 traversal and during zone 1 to 2 traversal were used to decode future movements (go to zone 4 vs. 5) and to decode the previously visited location (came from 4 vs. 5). For each spike, the 2-dimensional coordinates were projected onto a straight-line connecting zone 2 to 3 (or 1 to 2). Then, the distance between the projected point and zone 2 was calculated. Next, for each trial, dìerent spatial windows were applied incrementally, one decoding run at a time, across a range of spatial windows for the gap between two zones and discretized into equal-width bins. The spike numbers were counted for each bin to form the predictor of the logistic regression. Finally, each trial was labeled with the actual zone (zone 4 or 5) the rat reached at the end of that trial.

To decode the signals for previous memories, the data collection procedures were similar except that the label was marked as the zone it ‘Came From’ (zone 4 or 5). Then, the logistic regression model was adopted to build a binary classifier. A leave-one-out technique was used to evaluate model performances. That is, given the collection of spike counts of all trials, each trial was considered as the test data one trial at a time, and all remaining trials were the training data to build the logistic model. Finally, the test accuracy was obtained as the percentage of the correct prediction of all trials. In total, 29 behavior sessions were analyzed (29 for PC from rat 1-5, and 22 for hippocampus from rat 2-5).

### Temporal Neural Decoding of ‘Came From’ Location and ‘Go To’ Location

A binary logistic regression was applied to decode the rat’s future behavior and encoding of the past spatial location. The approach was similar to spatial decoding, except that due to dìerences in the time rats took to run the segments (Extended Data Fig. 1), all movements and spike times were standardized by dividing each time value by 5 s (a value that is longer than most running durations) for segments 1-2 and 2-3 separately. Next, for each trial, dìerent time windows were applied incrementally, one run at a time, across a range of time windows for the gap between two zones and discretized into equal-width bins. The spike numbers were counted for each bin from recorded tetrodes to form the predictor of the logistic regression. ‘Go To’ and ‘Came From’ decoding performance was assessed as for spatial decoding. In total, 29 behavior sessions were analyzed (29 for PC from rats 1-5, and 22 for hippocampus from rats 2-5).

### Route-Centered Analysis

As with spatial decoding, the rats’ trajectory was projected onto straight lines between each pair of zones. We split paths 4-1-2-3 and 5-1-2-3, and calculated the firing rates for each linearized path, averaging across runs through each path. Finally, the firing rate was converted to a z-score based on mean firing rates of the whole sequence. Correlations between the two paths were calculated for each segment to determine how similar (or dìerent) the firing rate by position was for each path.

### Place Cells

Place cells were defined by spatial information and coherence. Spatial information measures how much information a neuron’s firing conveys about the animal’s position. As described on Skaggs et al.^65^, the formula is:

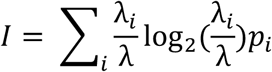

where λ_i_ is the firing rate in the i^th^ spatial bin, λ is the overall mean firing rate, p_i_ is the occupancy probability of the i^th^ spatial bin.

Coherence measures the local smoothness of the place cell’s firing rate map. It was calculated as the correlation between the firing rate of a bin and the average firing rate of its neighboring bins.

To further determine whether a neuron is a significant place cell, firing rates were shùled 1000 times, and corresponding spatial information and coherence were computed as control distributions. A cell was considered a place cell if its spatial information and coherence were both larger than 95th percentile of corresponding control distributions.

### Egocentric Goal Direction Analysis

Egocentric goal direction (also known as relative direction) was defined as the angle between the vector of the rat’s head direction and the vector from the rat to the goal. For each timestamp recorded, the corresponding relative direction was calculated. For each hippocampal single unit or PC MUA, a median resultant length (MRL) was computed by using CircStat Toolbox^66^. To test whether a single cell was significantly tuning, we shùled the cell’s head directions 1000 times such that the head directions were no longer associated with actual positions on the maze, as in Ormond and O’Keefe^17^. Single cells whose MRL was larger than 95 percentile of shùled distribution were considered significantly tuned to an egocentric goal direction. For PC MUA, due to the issue that shùle test was not able to handle high spiking frequencies (leading to false positives), we used dìerent criteria for PC MUA. We used a Rayleigh test combined with a between session stability test, which compared correlations of self-motions between two behavior sessions to 95 percentile of 1000-time shùled contributions, and change in peak vector did not change for more than 40 ° within one day (or between the first and second halves if there was only one behavior session), as in Wilber et al.^10^ MUA hat passed both tests was considered to have significant egocentric tuning.

For population vector fields, max spiking rates and the corresponding tuning direction of each significantly tuning single cells or MUA clusters were summed together. Then the mean directions of populations were computed by CircStat Toolbox.

### Trajectory Decoding

A Kalman Filter approach was applied to build a continuous model to predict the rat’s moving trajectories. The entire traversal for route 1-2-3-4 or 1-2-3-5 was evaluated. For each route, the time bin was fixed as one third of a second and the spike count was recorded for all tetrodes inside each bin. This formed the sequence of predictors. Next, the coordinates and velocities were computed on the x-and y-axes for each bin. This formed the sequence of responses. A conventional Kalman Filter was built on the sequence of predictors and response. During the training process, the leave-one-out technique was adopted to subsequently treat each trial as the test data, and the remaining trials as the training data. The correlation between the true and predicted trajectories was used to evaluate the model performance.

### Self-Motion Analysis

Position and head direction data were utilized to map the self-motion reference frame for each MUA cluster. For these analyses, position data were interpolated and smoothed by convolution with a Hamming window that was 1 s long. In addition, head direction data gaps < 1 s were transformed using directional cosines for interpolation using the interp1 function in MATLAB, then transformed back to polar coordinates^67^. Next, head angular velocity, linear velocity and MUA activity rate were calculated for each video frame using a 100 ms sliding window. Finally, the occupancy and number of MUA spikes for each 3 cm/s by 20°/s bin were calculated and converted to firing rate for each bin with > 0.5 s of occupancy. For illustrative purposes (not analyses), the self-motion activity rate maps were smoothed by convolving with a Gaussian function (σ = 1)^10,41^. The self-motion colormaps represented a range of MUA activity rates from 0 to the maximum. No adjustments were made to the standard, evenly spaced colormap. MUA clusters were classified as having a preferred self-motion state if the common points with sùicient occupancy (> 0.5 s) from the self-motion maps for the first and second daily session (or split one-half for two datasets with one daily session) were significantly positively correlated (p < 0.01). This was generally the most conservative criterion for self-motion cells of the three criteria reported by Whitlock et al.^41^.

### Overall Motion Prediction

We used the same approach as in our previous work^23^, which is to combine linear speed and angular speed (after normalization) into a single overall motion value, as is shown in the equation below:

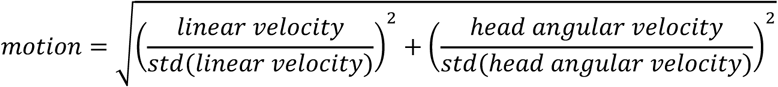

where std(⋅) denotes the standard deviation calculation. Note, that as a normalized measurement, the overall motion is unitless.

### Statistical Analysis

For statistics comparisons conducted on datasets from multiple rats, a mixed-effects model was used, and rat identity was included as a variable. Otherwise, paired t-tests were conducted on GraphPad Prism 10 or MATLAB. The mixed-effects model was adopted to examine the significance of the paired data. Since there exist multiple datasets originated from the same rat, the conventional t-test or ANOVA may not be an appropriate approach due to the violation of independence. Whenever there was paired data, the dìerence was first taken, and the mixed-effects model could determine if it dìers from zero. In addition, we conducted repeated measures two-way ANOVAs in case there were within-subject repeated measurements with respect to two factors. We examined the significance of each factor, as well as their interaction term. Except where noted otherwise, p < 0.05 was considered statistically significant. Sample sizes were not predetermined with statistical methods but were based on the standard for the field. Statistical details of the experiment can be found in this section of the methods, and in some cases in the results and figure legends. N can represent the number of tetrodes, datasets, trials, or rats and is specified where the data are presented. All descriptive statistics are presented as Mean ± Standard Error of Mean (SEM).

## Author Contributions

AAW and YZ designed the experiment, AAW, YZ, SCM conducted the experiment, AAW, YZ, XZ, SMR, LJD and WW analyzed the data, AAW, YZ, and XZ wrote the paper.

## Acknowledgements

This research was supported by grants from F32 MH099682, NIA K99/R00 AG049090, FL DOH 20A09, FL DOH 21K12 and R01 AG070094 to AAW.

## Notes

### Competing Interest Statement

The authors have declared no competing interest.

